# TEDDY: An integrative workflow for TE-chimeric isoform reconstruction and systematic characterization of TE-dependent transcriptional regulation

**DOI:** 10.64898/2026.08.22.746432

**Authors:** Yihan Xiao, Lu Shen, Cizhong Jiang, Yanping Zhang, Yuwei Liang, Jiqing Yin, Hong Wang, Jiejun Shi, Rongrong Le, Shaorong Gao

## Abstract

Transposable elements (TEs) substantially expand transcriptional regulation and complexity. Despite advances in sequencing technologies, the systematic interrogation of TE-dependent transcripts across diverse samples and platforms remains limited. Here we introduce TEDDY, a computational workflow for large-scale reconstruction, quantification and regulatory inference of TE-dependent isoforms. Using TEDDY, we resolved the landscape of TE-dependent isoforms in mammalian preimplantation development, revealing a conserved pattern of stage-specific isoform generation via species-specific TE exonization. We reconstruct a TE-dependent regulatory network underlying pluripotent-to-totipotent transition wherein key transcription factors are driven by TE-derived promoters and validated experimentally. Within this network, *Arid3a*, a novel regulator nominated by TEDDY, was functionally validated as essential for totipotency establishment and early development. Further application of TEDDY to hepatocellular carcinoma identified recurrence-associated prognostic isoforms, underscoring their clinical relevance. Benchmarking establishes TEDDY’s accuracy and efficiency with unique capabilities in full-length isoform recovery, cross-sample and -platform analysis, visualization, and TF–TE–gene network reconstruction, making TEDDY applicable across diverse biological contexts.

## INTRODUCTION

Transposable elements (TEs) make up a substantial fraction of mammalian genomes and influence gene expression through diverse mechanisms^1–5^. TEs can provide regulatory sequences, such as promoters, enhancers, and splice sites, thereby contributing to context-dependent gene regulation^6–12^. Beyond these cis-regulatory roles, TE sequences are frequently incorporated into host transcripts, resulting in chimeric TE–gene isoforms, which can confer post-transcriptional regulation or generate novel protein variants with distinct biological functions^13–16^. Investigating the functional importance of chimeric TE–gene transcripts has provided new insights into the molecular mechanisms of diverse biological processes in both physiological and pathological contexts^14–18^. For instance, a number of TE-chimeric transcripts have been detected during mammalian zygotic genome activation (ZGA)^14,19–21^. In innate immunity, TE exonization constitutes a core regulatory mechanism, exemplified by the generation of decoy receptor isoforms that potently inhibit inflammatory signaling^22^. Yet our current knowledge remains fragmented, built upon isolated examples rather than a systematic view. Therefore, to better understand the roles of TEs in gene transcription and evolution, it is essential to characterize TE-dependent transcripts and associated regulatory events in a comprehensive and systematic manner.

The development of RNA sequencing (RNA-seq) technology has opened new avenues for detecting TE-derived transcripts at a transcriptome-wide level. Currently, long-read platforms provide full-length transcript coverage, but their higher error rate and lower throughput limit quantification, especially in low-input samples^23,24^. Besides, raw long-read output must first be collapsed into a non-redundant isoform catalog and then harmonized across samples. Conversely, short-read RNA-seq offers cost-effective, high-depth coverage and additionally benefits from extensive public datasets. Therefore, computational tools are needed to fully leverage the advantages of diverse sequencing technologies for comprehensive transcriptome analysis. In particular, extracting biologically meaningful TE-dependent isoform events poses a significant computational challenge. This, in turn, calls for integrative, cross-platform computational frameworks.

In recent years, many computational methods have been developed to analyze TE-associated transcription, ranging from locus- or subfamily-level TE expression quantification to TE-initiated transcript detection, TE exonization analysis, and broader workflow-oriented approaches^15,25–30^. However, the available methods differ substantially in analytical scope and have distinct methodological limitations, as detailed in Supplementary Note 1 and Supplementary Table 1.^26,30^ With current methods, detecting isolated chimeric junctions without isoform-resolved reconstruction provides limited insight into full transcript structures and their dynamic switching. Moreover, the lack of a unified framework for cross-sample and cross-platform transcriptome integration makes it difficult to systematically identify context-specific TE-dependent isoforms.

Therefore, we have developed TEDDY, a comprehensive and scalable pipeline that can uniquely leverage both short- and long-read RNA-seq data to reconstruct, quantify, and characterize TE-dependent transcripts within a coherent framework. Solid benchmarking demonstrates that TEDDY achieves high accuracy, sensitivity, and speed against existing methods, with unique capabilities for full-length isoform recovery and robust cross-sample, cross-platform compatibility. We employed TEDDY to systematically investigate the role of TE-dependent transcripts across biological scales. TEDDY constructed a dynamic atlas of TE-dependent transcripts across stages of mouse preimplantation development from integrated multi-source data. Based on isoform-resolved analysis, TEDDY identified TE exonization events and TE-derived promoters that generate stage-specific isoforms during early development. Focusing on totipotency-associated candidates, we further validated their transcript structures by RT-PCR and Sanger sequencing, followed by low-input long-read sequencing. We next leveraged TEDDY to dissect the totipotency transcriptional network by jointly analyzing *in vitro* and *in vivo* models, uncovering TE-driven isoform-based regulatory mechanisms. *Arid3a*, nominated by TEDDY as a candidate regulator of TE-chimeric transcripts during *in vitro* totipotency establishment, was functionally validated as essential for early development. We then extended these insights to human embryogenesis, revealing a conserved strategy to generate stage-specific gene isoforms via species-specific TE exonization. Finally, application of TEDDY to multiple hepatocellular carcinoma cohorts identified recurrent TE-dependent isoform changes with prognostic associations, demonstrating its utility for large-scale analysis. Collectively, our work establishes TEDDY as a foundational resource and provides a unified, multi-scale perspective on TE-driven transcriptional dynamics.

## RESULTS

Our newly developed framework, TEDDY, demonstrates improvements in the analysis of TE-chimeric events, along with gains in computational efficiency and several additional features (Supplementary Note 1 and Supplementary Table 1). TEDDY is an R-integrated workflow that detects, reconstructs, and quantifies TE-chimeric transcripts at both bin- and transcript-level, with built-in visualization and tools to trace putative regulatory links between TE-derived promoters, transcription factors, and target genes (Fig. 1A). In this framework, TE-chimeric transcripts are structurally defined as mature isoforms containing TE-derived sequences, while TE-initiated transcripts represent the subset originating from a TE-derived 5’ end. TEDDY then links these structure-based features to expression and condition-specific exon usage to identify TE-dependent isoform events. Thus, TEDDY facilitates the discovery of alternative splicing driven by TEs and the reconstruction of TE-centric regulatory networks. Following performance validation in the benchmark (see Methods), we demonstrate TEDDY’s utility in four systems where TEs are known to be critically active: from early development—spanning mouse pre-implantation embryos and a 2C-like system—to human early embryos and disease, specifically hepatocellular carcinoma (HCC).

**Fig. 1.**
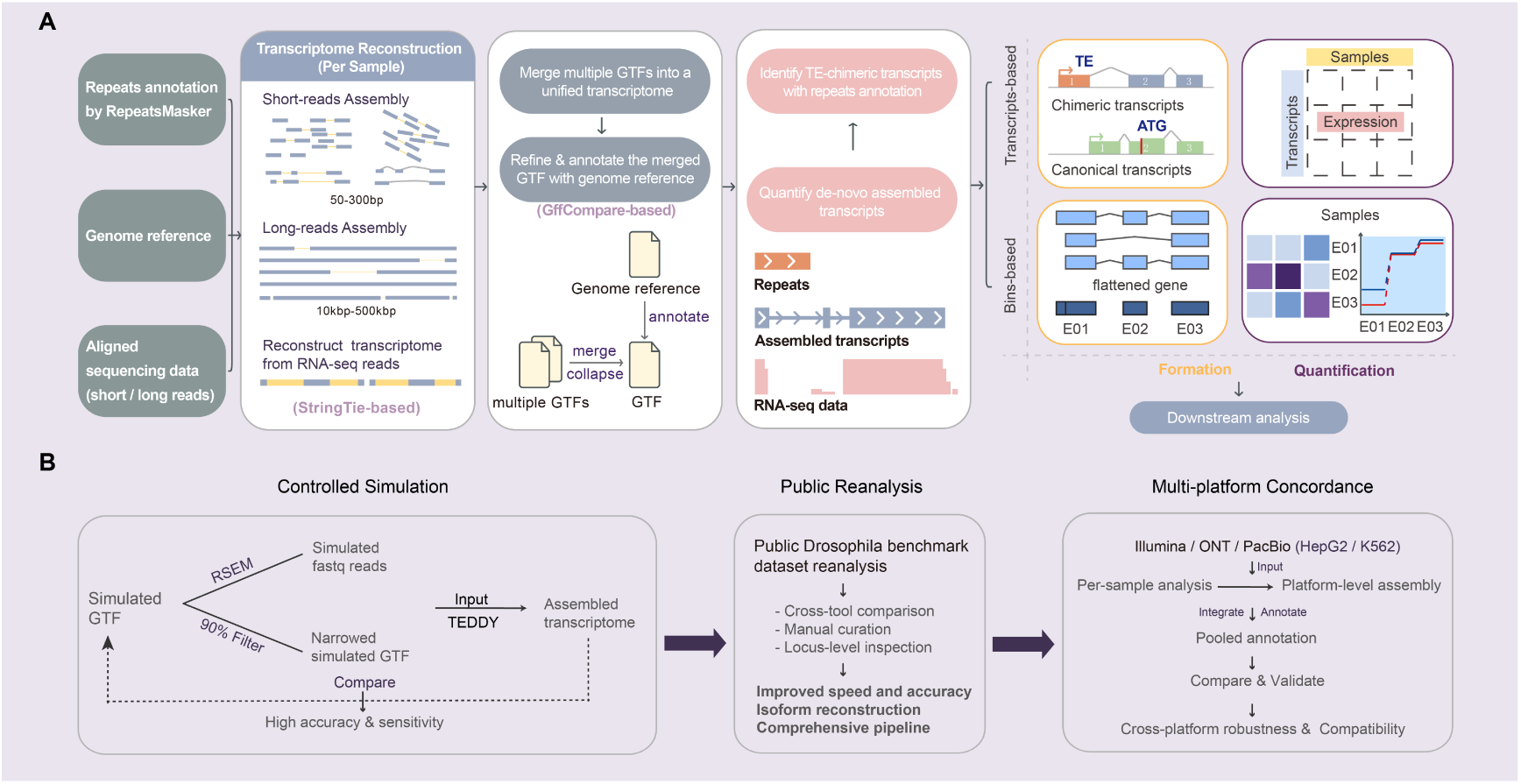
Overview of the TEDDY Workflow and Benchmarking Strategy. **A**, Analysis workflow. The method begins with transcript assembly to create transcriptome. The initial input data include repeat annotation, the reference genome in GTF format, and aligned sequencing data in BAM format. TEDDY predicts TE-chimeric transcripts through transcript assembly to reconstruct a unified multi-sample reference genome. TEDDY provides the structure and quantification information of TE-chimeric transcript at the transcript level (top right) and bin level (middle right). At the transcript level, TE sequences that are spliced into mRNA to form TE-chimeric transcripts can be captured; at the bin level, flattening genes into disjoint exonic bins provides a comprehensive view of exon usage across various isoforms. Together, these outputs support downstream analyses, such as visualization and investigation of alternative isoforms, as well as specific TE-chimeric events and associated regulatory mechanisms. **B**, Benchmarking strategy. Simulated GTFs, derived from real transcript annotations, served as the ground truth. These GTFs were used to generate simulated reads with RSEM and to construct a partial reference for benchmarking. The simulated reads and this partial reference were then used as inputs to evaluate the performance of TEDDY and comparator tools.

### In silico benchmarking demonstrates accurate detection of TE-chimeric loci

In silico simulations were used to quantify detection accuracy, sequencing-depth dependence, and computational characteristics under controlled conditions (see Methods and Fig. 1B, left). This approach was necessary because no comprehensive gold standard exists for TE-chimeric isoforms and the true transcript structure in real samples is generally unknown.

As noted above, the benchmarked tools differ in analytical scope and structural resolution. Some methods primarily detect TE-initiated candidates, while others report breakpoint-, junction-, or event-level outputs, and their built-in criteria defining TE– chimeric events and output formats are often not directly tunable. To enable a fair comparison, we benchmarked TEDDY against representative workflows for detecting TE-chimeric transcripts and TE-gene events, using identical aligned reads and partial-reference inputs. Performance evaluation was harmonized at the TE–chimeric locus level, and compatible ground-truth sets were matched to each tool’s intended scope. Performance was evaluated using precision, recall, and F1 score, based on true positives (TP), false positives (FP), and false negatives (FN) (Methods and Supplementary Table 1). TEDDY achieved high precision across sequencing depths, with recall and F1 increasing markedly as read depth increased (Fig. 2A). The other evaluated tools showed lower overall F1 scores, reflecting either limited recall for compatible TE-chimeric loci or reduced precision from broader event-level predictions.

**Fig. 2.**
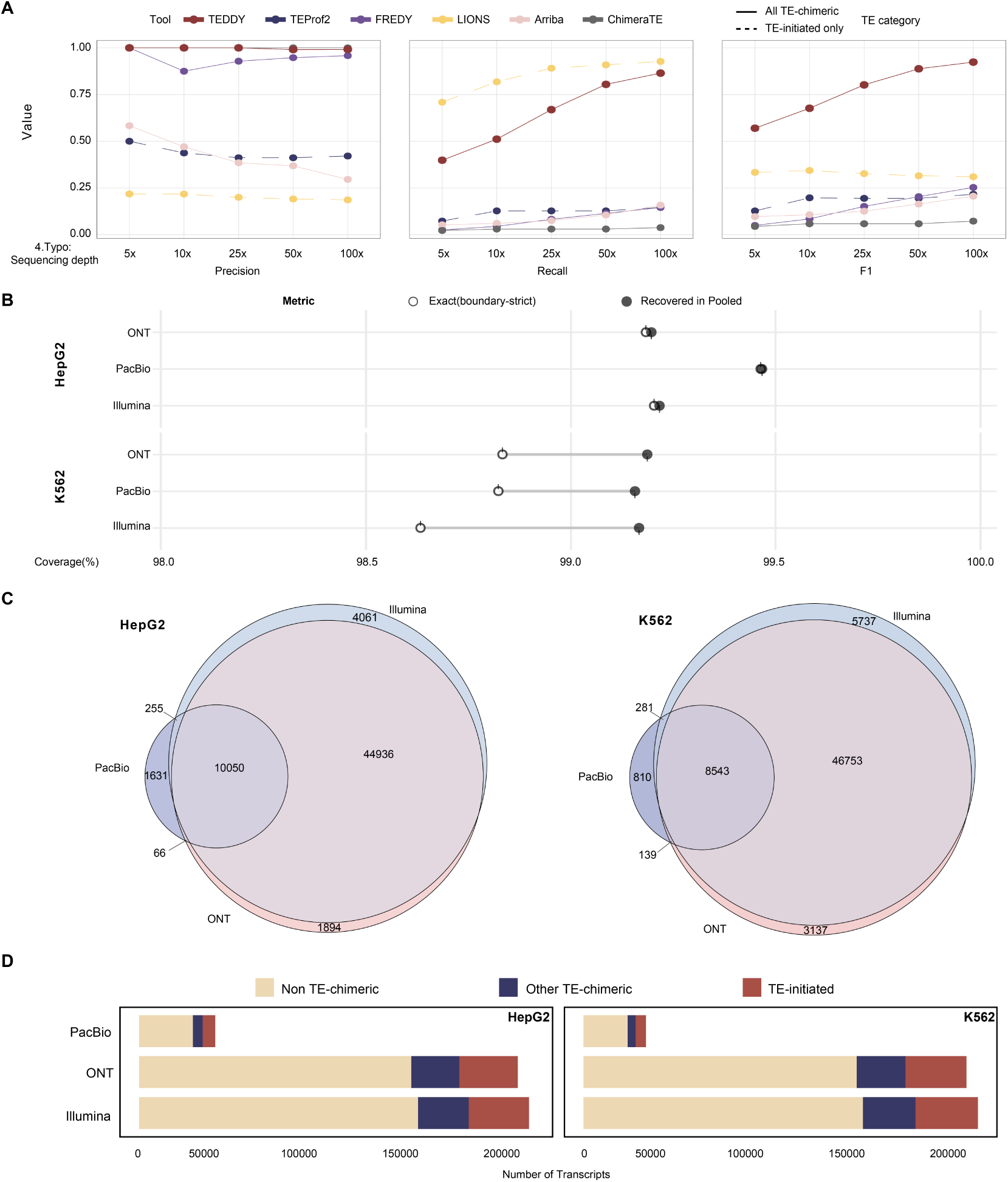
Comprehensive Benchmark Demonstrates TEDDY’s Accuracy, Resolution, and Versatility. **A**, Performance on simulated RNA-seq data across 5×–100× coverage. TEDDY (dark red) was evaluated using a strict isoform-level criterion, while comparison tools (LIONS, Arriba, ChimeraTE) were assessed using a permissive junction-overlap rule. TEDDY maintains near-perfect precision at all depths, while recall and F1 increase with coverage. **B**, Cross-platform integration accuracy. The pooled transcriptome reference generated by TEDDY’s merge process recovers ∼99% of TE-chimeric transcripts that were originally detected in individual platform-specific analyses (Illumina, ONT, PacBio). This high recovery rate with minimal loss (∼1%) validates the high fidelity of TEDDY’s integration across diverse sequencing technologies. **C**, Overlap of TE-chimeric transcripts across platforms by cell line. Venn diagrams show the isoforms detected by Illumina, ONT, and PacBio in the HepG2 (left) and K562 (right) cell lines. **D**, Distribution of transcript categories by platform. Stacked bars show the proportions of transcripts in three mutually exclusive categories: TE-initiated, Other TE-chimeric, and Non TE-chimeric, as defined by the location of TE sequences.

As illustrated by an example locus (Fig. S1A), increasing sequencing depth from 5× to 25× and 50× also improved the accuracy of local isoform reconstruction, reducing an ambiguous assembly isoform that contained an extra exon. These results demonstrate that sequencing depth is important for both TE-chimeric locus detection and local isoform-structure reconstruction. Nevertheless, these benchmarks represent controlled conditions and may not capture the full complexity of endogenous transcriptomes. We therefore complemented simulations with real-data and cross-platform evaluations below.

### Reanalysis of a non-mammalian dataset confirms accuracy and speed

We further validated TEDDY on a public *Drosophila* test subset released with ChimeraTE^31^, to avoid dataset-selection bias and to demonstrate that TEDDY generalizes beyond the mammalian datasets used elsewhere in this study (Fig. 1B, middle).

TEDDY and ChimeraTE shared four high-confidence TE-chimeric events, with one unique call reported by each tool (Fig. S1B). Manual inspection shows that TEDDY reconstructed a novel isoform at the *FBgn0031188* locus with strong coverage support (Fig. S1C), whereas ChimeraTE assigned this as an S2-chimeric event on the opposite strand, indicating a false positive. Conversely, at the *FBgn0262731* locus, TEDDY predicted an FB-chimeric isoform supported by strand-concordant coverage and junction-spanning reads, suggesting a ChimeraTE false negative. Runtime and memory measurements are provided in Supplementary Table 1 and Supplementary Note 1. Beyond event-level detection, TEDDY further reconstructs full-length isoform structures, including exon composition and splice-junction coordinates.

### Cross-platform evaluation demonstrates reproducibility and integration

To evaluate TEDDY’s robustness across sequencing technologies, we applied the pipeline to matched short- and long-reads datasets from the NeX project^32^, generated on Illumina, Oxford Nanopore, and PacBio platforms for K562 and HepG2 cell lines, which exhibit frequent fusion-like transcriptional events and detectable TE activity (see Methods and Fig. 1B, right).

Using TEDDY’s merge-and-annotate workflow to combine platform-level assembled GTFs into a pooled annotation for each cell line, we could assess whether platform-specific isoforms assigned arbitrary transcript IDs were recovered or matched in assemblies from the other platforms.

We mapped platform-specific isoforms back to the pooled reference using a permissive matching rule that tolerates small splice-site shifts. The high recovery (∼99%) and correspondingly low merge loss (∼1%) indicate that the vast majority of platform-specific isoforms can be recovered in the pooled reference (Fig. 2B). Requiring exact exon-chain concordance only marginally reduced recovery, indicating that TEDDY’s multi-sample, multi-platform merge accurately reconstructs exon composition and boundaries while producing few spurious, near-duplicate novel isoforms (Fig. 2B). Consistently, downsampling of the real K562 and HepG2 Illumina and ONT cDNA datasets showed high exact recovery of full-depth TE-chimeric isoforms within each platform. High recovery was also observed for TE-chimeric isoforms jointly supported by Illumina and ONT in the cross-platform analysis. For these exact-matched, Illumina–ONT-supported TE-chimeric isoforms, FPKM estimates from independently reconstructed downsampled Illumina datasets showed increasing concordance with the corresponding full-depth estimates as sequencing depth increased, supporting the stability of TEDDY’s structure-coupled quantification (Supplementary Table 2).

Most TE-chimeric transcripts are reproducibly recovered by two or more technologies, while platform-unique calls represent a minority (Fig. 2C). PacBio yields fewer calls in our pooled set, likely due to the lower sample number and coverage typical of long-read sequencing. Nevertheless, per-platform composition shows broadly similar distributions across three classes: non-TE transcripts, TE-initiated transcripts, and other TE-chimeric transcripts. In the two human cell lines studied, approximately 55% of TE-chimeric transcripts are TE-initiated, suggesting that TEs provide the transcription start sites for a substantial fraction of these TE-chimeric events (Fig. 2D). The broadly consistent patterns support reproducible biological interpretation across platforms and samples, while the integration of short- and long-read data enables more comprehensive capture of TE-chimeric transcript diversity across platforms, even under varying sequencing depths and data quality.

Although long-read sequencing provides more direct transcript-level evidence, long-read data remain far less prevalent than short-read sequencing. Our benchmark shows that TEDDY performs well on short-read data alone, enabling the reuse of large-scale short-read collections to detect and interpret TE-chimeric events. By leveraging StringTie’s ability to integrate short- and long-read sequencing data, TEDDY identified a cost-effective workflow readily compatible with multi-omics integration.

### The landscape of TE-chimeric transcription in mouse early embryos

There is a growing appreciation for the functional importance of TEs in early embryogenesis, as demonstrated by previous findings^10,16,19,33^. In addition to acting as regulatory elements, TE sequences can be incorporated into gene transcripts (TE–gene chimeric transcripts), greatly expanding the complexity of the host transcriptome^14^. Therefore, we utilized TEDDY to characterize transcriptome-wide TE-derived chimeric transcripts (termed TE-chimeric transcripts hereafter) and explore the underlying mechanisms. To assess TEDDY’s performance in analyzing large-scale datasets, we initiated our study with a specific focus on mouse preimplantation embryos. Approximately 12,000 TE-chimeric transcripts exhibited stage-specific dynamics and nine prominent expression patterns emerged, each corresponding to a distinct temporal profile across early embryonic stages based on the TEDDY-derived expression matrix (see Methods and Supplementary Table 3). We found that oocytes harbored the greatest number of TE-chimeric transcripts among all stages, of which a large portion was shared between oocytes and 2-cell embryos. Approximately half of the TE-chimeric transcripts detected in 2-cell embryos were specifically activated during zygotic genome activation (ZGA) and were rapidly downregulated from the 4-cell stage onward (Fig. 3A). TE-chimeric transcripts exhibited low expression levels from 8-cell to morula stages, whereas a large number of TE-chimeric transcripts were specifically activated at the blastocyst stage. Together, TE-chimeric transcripts exhibited highly dynamic and stage-specific expression patterns in early embryogenesis (Fig. 3A,B).

**Fig. 3.**
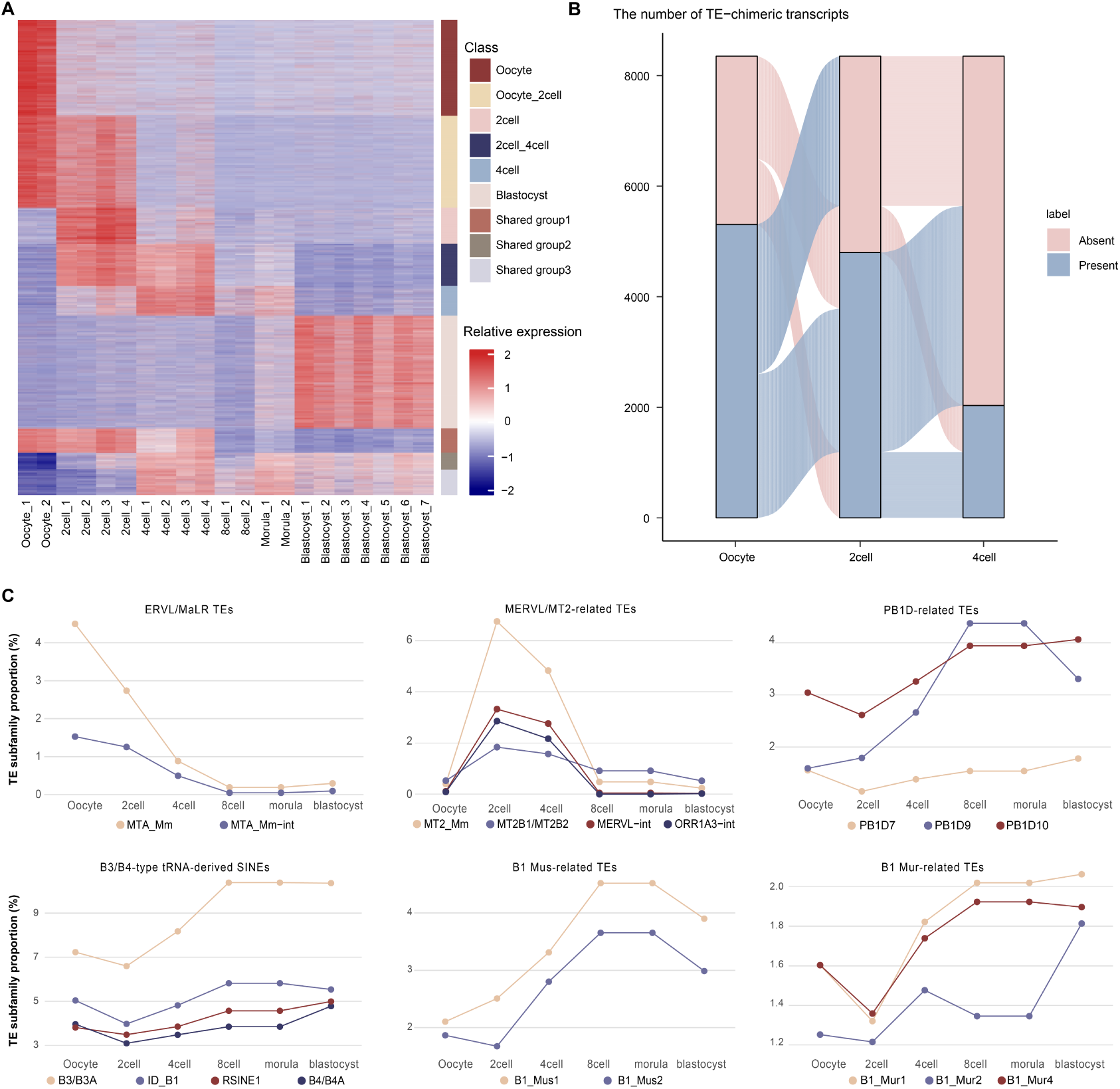
Characterization of TE-chimeric Transcripts in Mouse Preimplantation Development. **A**, Heat map generated from cluster analysis of TE-chimeric transcripts in mouse preimplantation embryos identified by TEDDY based on the expression patterns. Each row represents the Z score of log2-transformed FPKM (FPKM, fragments per kilobase of transcript per million) values. **B**, Alluvial plots showing the global dynamics of TE-chimeric transcripts across the oocyte, 2-cell, and 4-cell stages. **C**, Line plots showing the developmental dynamics of representative TE subfamilies associated with TE-chimeric transcripts in the expression clusters identified in 3A.

Next, we analyzed the contribution of TE subfamilies to the formation of TE-chimeric transcripts (Fig. 3C and Supplementary Table 3). ERV-derived elements consist of internal retroviral-like sequences flanked by LTRs, and solo LTRs can arise during genome evolution^34,35^. We therefore grouped related internal and LTR/solo-LTR elements together, including MERVL-int with its associated MT2 LTRs, and MTA_Mm-int with MTA_Mm. We found that MTA-related elements were enriched in early development, specifically at the oocyte stage (Fig. 3C) and within oocyte-associated expression groups (Supplementary Table 3), and declined during subsequent development. In contrast, MT2/MERVL-related elements, including MT2_Mm, MERVL-int, MT2B1/MT2B2, and ORR1A3-int, were most prominent in the 2-cell stage. Several SINE elements, including B3/B3A, B1_Mus1/B1_Mus2, ID_B1, RSINE1, and B4/B4A, were more broadly represented from the 4-cell stage onward and in later shared expression groups. Together, the composition of TEs incorporated into chimeric transcripts differed across clusters, reflecting the temporal dynamics of TE utilization during early development.

We further utilized TEDDY to perform a detailed analysis on the structure of TE-chimeric transcripts. We first focused on previously identified gene loci generating TE-chimeric transcripts in oocytes to evaluate the accuracy of TEDDY^20^. TEDDY detected an MTB-initiated truncated isoform and a non-chimeric isoform of *Spin1* in oocytes (Fig. S2A,B), which was experimentally validated in previous studies^20^. Similarly, the previously validated chimeric isoform of *Vdac2*, which was predominantly expressed in oocytes and initiated from an MTC element ∼6.5 kb upstream, was also captured (Fig. S2C,D)^20^. For *Dnajc11*, TEDDY recovered the reported MTC-initiated TE-chimeric isoform (ENSMUST00000142198.6) and identified previously unannotated TE-initiated isoforms with distinct exon usage (for example, MSTRG.84207.16) (Fig. S2E,F).

### Characterization of TE-dependent transcripts in 2-cell embryos and 2C-like cells

Recent studies indicate that TEs play important functional roles in zygotic genome activation (ZGA) and the establishment of totipotency^8,19,33,36^. It has been shown that TE-derived cis-regulatory elements play an important role in controlling gene expression in early embryos^10,16,37^. In particular, MERVL, one of the most abundantly expressed TEs in 2-cell embryos, has been shown to serve as promoters to initiate transcription of totipotency-related genes^37^. However, it remains unclear whether TE sequences could be incorporated into gene transcripts to generate alternative untranslated regions (UTRs) or open reading frames (ORFs) beyond their cis-regulatory roles in totipotent cells. To investigate this question, we examined TE-dependent transcripts in 2-cell embryos and totipotent-like stem cells (2C-like cells)^38^. 2C-like cells are a transient subpopulation of mouse embryonic stem cells exhibiting transcriptional and chromatin features similar to those of the 2-cell embryos, including activation of MERVL and *Zscan4*, and are widely used as a highly valuable *in vitro* model to study totipotency^38^. We therefore applied TEDDY to analyze TE-dependent transcriptional events in 2-cell embryos and 2C-like cells.

After characterizing expression patterns across ESCs, 2C-like cells and *in vivo* preimplantation embryos (Fig. S3), we conducted a comprehensive analysis of TE-dependent transcriptional events. We found 4,214 transcripts that were induced in 2C-like cells compared with ESCs, and 2,995 of these were also expressed in 2-cell embryos *in vivo* (hereafter referred to as 2C-like transcripts), defining a shared 2C-like transcriptional program (Figs. 4A and S4). Notably, 1,651 of the shared transcripts were TE-chimeric isoforms detected by TEDDY, indicating a strong contribution of TE-chimeric transcripts to the 2C-like state.

**Fig. 4.**
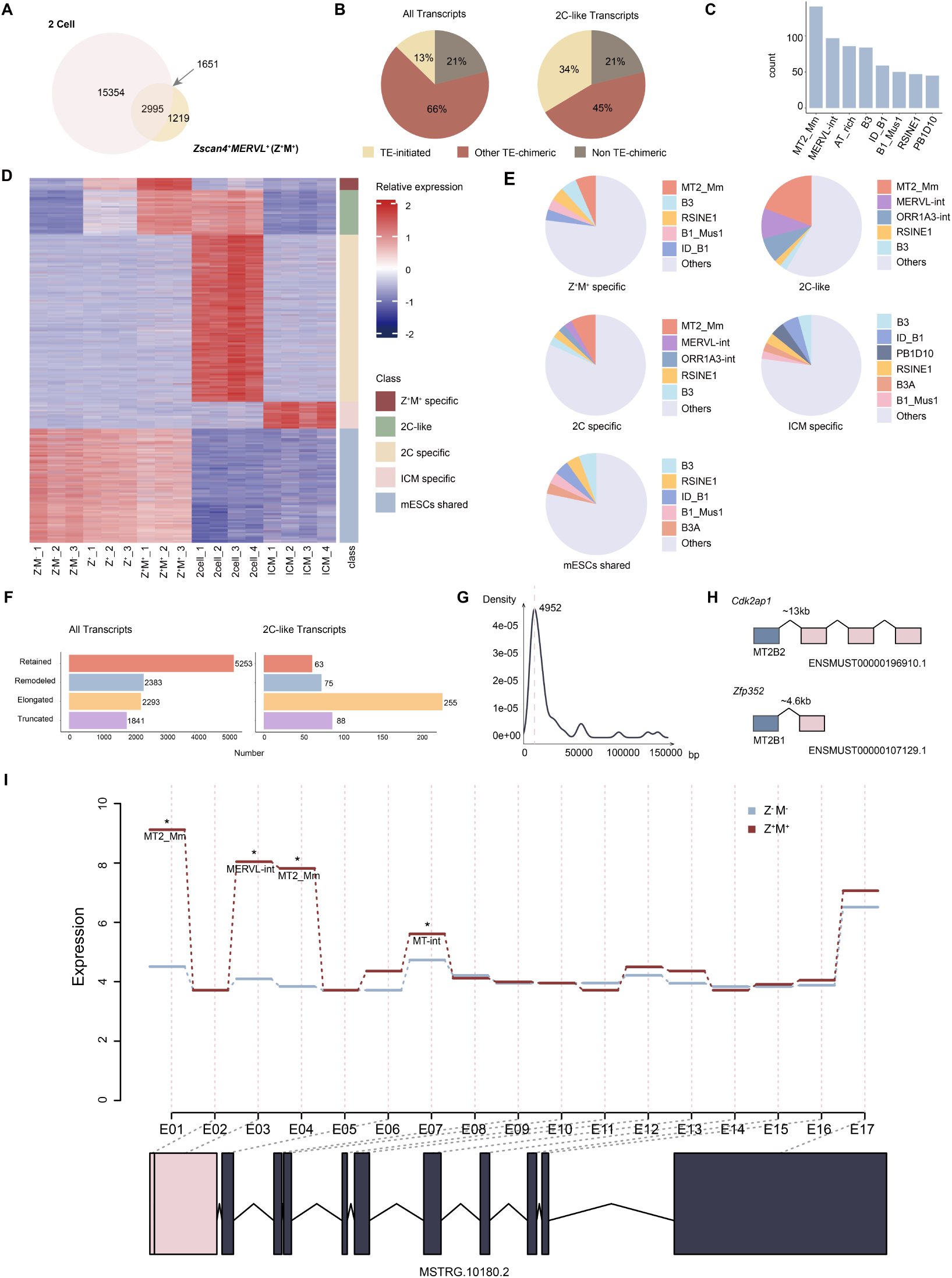
Identification of TE-chimeric Transcripts in 2-cell Embryos and 2C-like Cells. **A**, Venn diagram shows 2995 transcripts shared between 2-cell embryos and 2C-like cells (among which 1651 transcripts are TE-chimeric isoforms). These transcripts were identified through DESeq2 analysis, which displayed significant differential expression changes between 2C-like cells and ESCs (log2 expression fold change greater than 1 and an adjusted *P*-value less than 0.01). **B**, Pie charts showing the proportions of TE-initiated chimeric transcripts, TE-chimeric transcripts that are not initiated from TEs and non-chimeric transcripts in preimplantation embryos (left) and 2C-like cells (right), indicating a higher proportion of TE-chimeric transcripts in 2C-like state. **C**, Bar plot showing the top enriched TE subfamilies among the TE-initiated chimeric transcripts identified as significant by TEDDY’s bin-level differential analysis in 2C-like cells (adjusted *P* < 0.01); MERVL (MT2_Mm and MERVL-int) is the predominant family. **D**, Heat map generated from cluster analysis of TE-initiated transcripts in ESCs, MERVL^+^/*Zscan4*^+^ cells, *Zscan4*^+^ cells, 2-cell embryos and blastocyst identified by TEDDY based on the expression patterns. Five groups of TE-chimeric transcripts were categorized. Each row represents the Z score of log2-transformed FPKM values. Z^-^M^-^: ESCs; Z^+^: *Zscan4*^+^ cells (intermediate cell state in the transition between ESCs and 2C-like cells); Z^+^M^+^: 2C-like cells. **E**, Pie charts showing the proportion of TE subfamilies associated with TE-initiated transcripts in the different groups identified in 4D. **F**, Structural categories of TE-initiated transcripts. Bar plots show the number of transcripts classified into four structural categories (Elongated, Truncated, Retained, Remodeled). Left: all TE-initiated transcripts identified in preimplantation embryos. Right: TE-initiated transcripts shared between 2C-like cells and 2-cell embryos (2C-like transcripts) **G**, Density plot shows the genomic distances between TEs and gene exons to generate TE-initiated transcripts. **H,** Schematic representation for structures of MT2B2-initiated *Cdk2ap1* TE-chimeric isoform (top) and the TE-chimeric isoform of 2C-like gene *Zfp352* (bottom) generated by TEDDY. The genomic distances between TEs (grey) and gene exon (pink) are indicated. **I**, Step plot showing exon usage and the normalized RNA-seq read counts of individual exons of *Fam172a* in 2C-like cells (red) and ESCs (blue). The x-axis displays collapsed exon bins across the gene range. TEs with the potential to drive gene expression predicted by TEDDY in 2C-like cells are marked with asterisks. The bottom diagram illustrates the structure of the TE-initiated isoform of *Fam172a*, with dotted lines linking its included exons.

Globally, TE-chimeric isoforms exhibited markedly elevated expression in 2C-like cells, mirroring their activation in the *in vivo* 2-cell stage (Fig. S4A). Among these TE-chimeric transcripts, TE-initiated isoforms constituted a major proportion, and this proportion was even higher in 2C-like transcripts (Fig. 4B). MERVL-associated elements were highly prevalent in these 2C-like TE-initiated transcripts (Fig. S4B). These observations suggest that TE sequences are not only incorporated into transcripts but may also function as cis-regulatory elements, specifically as alternative promoters to activate 2C-like transcription.

Thus, using TEDDY’s built-in generalized linear model (GLM) on bin-level expression counts (see Methods), we assessed locus-specific changes in TE-bin usage to evaluate their putative cis-regulatory contributions. Candidate TE-chimeric transcripts supported by significant TE-bin usage (adjusted *P* < 0.01) were selected for downstream analysis. A summary of this analysis is presented in Table 1, with output-field definitions provided in the table legend. The analysis identified MERVL (MT2_Mm and MERVL-int) as the predominant regulatory influence driving the TE-chimeric transcriptome during the 2C-like transition (Fig. 4C).

**Table 1.** Representative results of TE influence on transcript dynamics in 2C-like state transition.

| groupID | featureID | exonExpr | TEclass | dispersion | stat | pvalue | padj | geneName |
| --- | --- | --- | --- | --- | --- | --- | --- | --- |
| MSTRG.10180 | E001 | 268.1169 | MT2_Mm,MERVL-int | 0.01220744 | 347.5465688 | 1.45E-77 | 1.49E-74 | Fam172a |
| MSTRG.120 | E018 | 2.311576 | MT2_Mm | 0.058078026 | 12.14285955 | 0.000492762 | 0.012176256 | Sgk3 |
| ... | ... | ... | ... | ... | ... | ... | ... | ... |
| MSTRG.137 | E045 | 44.96797 | MT2_Mm | 0.0293428 | 53.44173655 | 2.66E-13 | 3.00E-11 | Prex2 |
| MSTRG.27328 | E018 | 306.9122 | MT2_Mm,B1_Mus1 | 0.008742988 | 276.0855823 | 5.35E-62 | 4.10E-59 | Nelfa |
| MSTRG.10348 | E008 | 700.9222 | MT2_Mm,MERVL-int,ORR1A3-int | 0.078472515 | 24.54336818 | 7.27E-07 | 3.93E-05 | Scamp1 |

Following a recent study showing that CRISPR interference of MT2_Mm in 2-cell embryos downregulates ZGA genes and arrests development^37^, we reanalyzed the expression changes of 2C-specific genes predicted by our algorithm to be driven by MT2_Mm/MERVL-int (Fig. S4C,D). As expected, repression of MT2_Mm caused a significant reduction in the expression levels of MT2-driven chimeric transcripts predicted by TEDDY (Fig. S4C,D), supporting TEDDY’s reliability to predict the potential of TEs to influence gene expression.

We next systematically catalogued TE-initiated transcripts by clustering their expression across 2-cell embryos, blastocysts, 2C-like cells (MERVL^+^/*Zscan4*^+^ cells), intermediate MERVL^-^/*Zscan4*^+^ cells and ESCs (See Methods). Five distinct clusters of TE-initiated transcripts were resolved, revealing stage-specific TE utilization patterns (Fig. 4D,E). MERVL (MT2_Mm, MERVL-int, ORR1A3-int) elements are more frequent sources of TE-initiated isoforms in both 2C-like cells and 2-cell embryos (Fig. 4E). In contrast, blastocyst-enriched TE-initiated isoforms were predominantly SINE-derived (B3, ID_B1, PB1D10; Fig. 4E). We next analyzed TE-initiated isoforms in *in vitro* 2C-like cells, alongside their *in vivo* counterparts. ∼600 were induced during the transition from ESCs to 2C-like cells (cluster “2C-like”), while ∼1,600 remained stable (cluster “ESCs-shared”; Fig. 4D). Of the TE-initiated transcripts upregulated in 2C-like cells versus ESCs, ∼78% were also expressed in *in vivo* 2-cell embryos (Fig. 4D), indicating that the 2C-like transition toward totipotency is strongly aligned with TE-dependent transcription. Collectively, 2-cell embryos and 2C-like cells share a distinct MERVL-driven TE-initiation transcriptional signature associated with totipotency.

While the role of TEs in the 2C-like transcriptome has been emphasized, the structures, splice sites, and full-length sequences of the resulting chimeric isoforms require resolution to clarify the mechanisms of these events in the 2C-like state. We therefore applied the core analysis module of TEDDY to delineate how TEs alter transcript structures. We compared TE-initiated isoforms with their non-chimeric counterparts in preimplantation embryos and 2C-like cells (Fig. 4F). This analysis identified a large number of novel and GENCODE-annotated chimeric transcripts. We then classified TE-initiated transcripts into four types (Fig. 4F): (1) Elongated: TEs located upstream of genes drive the production of elongated gene isoforms; (2) Truncated: TEs initiating transcripts downstream of all annotated transcript start sites drive the production of truncated gene isoforms; (3) Retained: TEs located within gene loci drive the generation of chimeric isoforms already annotated by GENCODE (e.g., *Dnajc11* chimeric isoform ENSMUST00000142198.6; Fig. S2E); (4) Remodeled: TEs located within gene loci lead to differential exon usage patterns (e.g., MSTRG.84207.20, a novel *Dnajc11* chimeric isoform detected by TEDDY; Fig. S2E). TEDDY identified numerous TE-initiated transcripts in mouse preimplantation embryos, comprising both novel (elongated, truncated or remodeled) and GENCODE-annotated (retained) ones (Fig. 4F, left). Of note, most of the TE-initiated transcripts shared by 2-cell embryos and 2C-like cells were unannotated, highlighting the need to reassess the TE-dependent isoforms during totipotency acquisition (Fig. 4F, right).

Prior work primarily centered on MT2-proximal 2-cell genes to probe TE enhancer and promoter activity, limiting the detection of distal TE–gene interactions^38–40^. Thus, we determined the genomic distance between TEs and the downstream gene exons to form chimeric splicing junctions. Although chimeric splicing events preferentially occurred across short genomic distance (∼5 kb), long distances up to 150 kb were also observed (Fig. 4G), a regime poorly captured by existing proximity-focused, junction- or soft-clipped–based methods. The MT2B2-driven isoform of *Cdk2ap1* promotes cell proliferation in early embryos, exhibiting an opposite function to its canonical counterpart^16^. This chimeric isoform is driven by an MT2B2-derived promoter located 8.2 kb upstream of *Cdk2ap1*^16^. This visualization illustrates TEDDY’s capacity to identify long-distance TE-chimeric events (Fig. 4H, top). Our structural information further indicates that the MT2B2-initiated *Cdk2ap1* isoform skips the classical exon 1 and generates a truncated isoform, as reported. In addition, we identified TE–gene splicing in the *Zfp352* gene locus, an important totipotency-associated transcription factor^41^, with an MT2B1-derived TSS ∼4.6 kb upstream of the proximal exon (Fig. 4H, bottom). Therefore, TEDDY is capable of resolving detailed transcript structures and alternative splicing events of chimeric TE–gene isoforms.

Beyond recapitulating known transcripts, TEDDY predicted numerous novel TE-chimeric gene isoforms in 2C-like cells. Manual inspection of these predictions using the IGV Genome Browser corroborated their plausibility, prompting experimental follow-up (Supplementary Table 4). We experimentally confirmed novel isoforms of key candidate 2C-like factors, including *Nelfa, Snai1*, *Pou6f2*, and *Lmx1a* via RT-PCR and Sanger sequencing (Fig. S5A and Supplementary Table 5). The maternal factor Nelfa, known as an essential transcriptional activator of *Dux* that drives the 2C-like state, serves as a compelling instance^42^. A previously unreported TE-initiated isoform of *Nelfa* was detected by TEDDY and experimentally validated (Fig. S5A,B). Locus-level analysis revealed that this MT2_Mm-initiated isoform is N-terminally truncated relative to its canonical non-chimeric counterpart (Fig. S5A,B). The expression level of the TE-chimeric isoform of *Nelfa* was much greater than that of the canonical non-chimeric isoforms in 2C-like cells (Fig. S5C). These experimentally validated isoforms encode transcription factors themselves, hinting that TE-derived promoters may directly rewire the transcriptional regulatory network in 2C-like cells.

Importantly, these isoforms were identified by TEDDY solely from the original short-read datasets using GENCODE M7 as the reference annotation. To assess the robustness of TEDDY to reference-annotation updates, we reran the analysis within the same mm10 coordinate framework using GENCODE M25 and found that the candidate structures were broadly retained across annotation versions. Notably, only 22 of the 570 novel isoforms (3.9%) were represented in the official GENCODE M25 catalog. Moreover, subsequently generated low-input Nanopore data from 2-cell embryos independently supported 553 of 570 novel TE-initiated isoforms (97.0%) at the TE-host junction level and 461 of 570 (80.9%) at the exon-chain level (Supplementary Table 4). Together, these orthogonal analyses support both the structural accuracy of TEDDY’s short-read-based predictions and the robustness to reference-annotation updates.

To further characterize complex splicing events and the resulting alternative isoforms, TEDDY integrates transcript-structure visualization with quantitative comparison of exon-bin usage across conditions and samples, as exemplified by *Fam172a* (Fig. 4I). The MT2_Mm-initiated chimeric isoform, specifically expressed in 2C-like cells compared to ESCs, together with its exon-usage pattern, is depicted. In summary, TEDDY enables comparative analysis of TE-associated transcript structures, exon usage, and expression across samples and conditions.

### Construction of TE-dependent transcriptional network underlying pluripotent-to-totipotent state transition

TEs provide abundant binding sites of transcription factors (TFs) to modulate gene expression in both pathological and physiological contexts^6,43^. Motivated by the high prevalence of TE-derived promoters in totipotent cells, we first performed an exploratory motif-enrichment analysis on TE genomic sequences associated with TEDDY-defined TE-derived promoters using HOMER findMotifs.pl^44^. This identified highly enriched motifs for established critical totipotency factors (Dux, Obox1)^45–48^, core pluripotency regulators (e.g., Prdm14, Nanog, Klf5), and newly implicated factors including Arid3a and bHLH family members, suggesting their potential role in activating MERVL-derived promoters of 2C-related genes (Fig. S6A). To prioritize these candidates, we examined their expression across ESCs (pluripotent stem cells), *Zscan4*⁺ cells (intermediate cells of 2C-like transition), and 2C-like cells (totipotent-like stem cells) and across early embryonic stages (Fig. S6B,C). These TFs exhibited significant expression changes, further indicating their potential regulatory function. Although the major players in the totipotency network, such as Dux and Obox1, have been reported to activate MERVL and 2C-like genes, their precise roles in the transcriptional logic of totipotency remain unclear^47,49–51^. To address this gap, we analyzed published Dux ChIP-seq data^51^ and found that Dux peaks frequently overlapped repetitive elements, with MT2_Mm and MERVL-int being the most prominent TE subfamilies (Fig. S7A,B). We observed recurrent Dux occupancy at TEDDY-defined TE-derived promoters, including multiple representative loci such as the totipotent regulator *Zfp352* (Figs. 5A and S7C,D; Table 2)^38,41^. TEDDY identified an upstream MT2B1 element bound by Dux driving the TE-chimeric isoform of *Zfp352*. Furthermore, we validated a novel, truncated TE-chimeric isoform of *Lmx1a* (as shown in Fig. S5A), whose expression is driven by an intronic MT2_Mm element with robust Dux occupancy (Fig. S7D). Collectively, these results support a direct role of Dux in the activation of TE-chimeric transcripts during 2C-like transition, across both annotated and TEDDY-reconstructed novel isoforms.

**Fig. 5.**
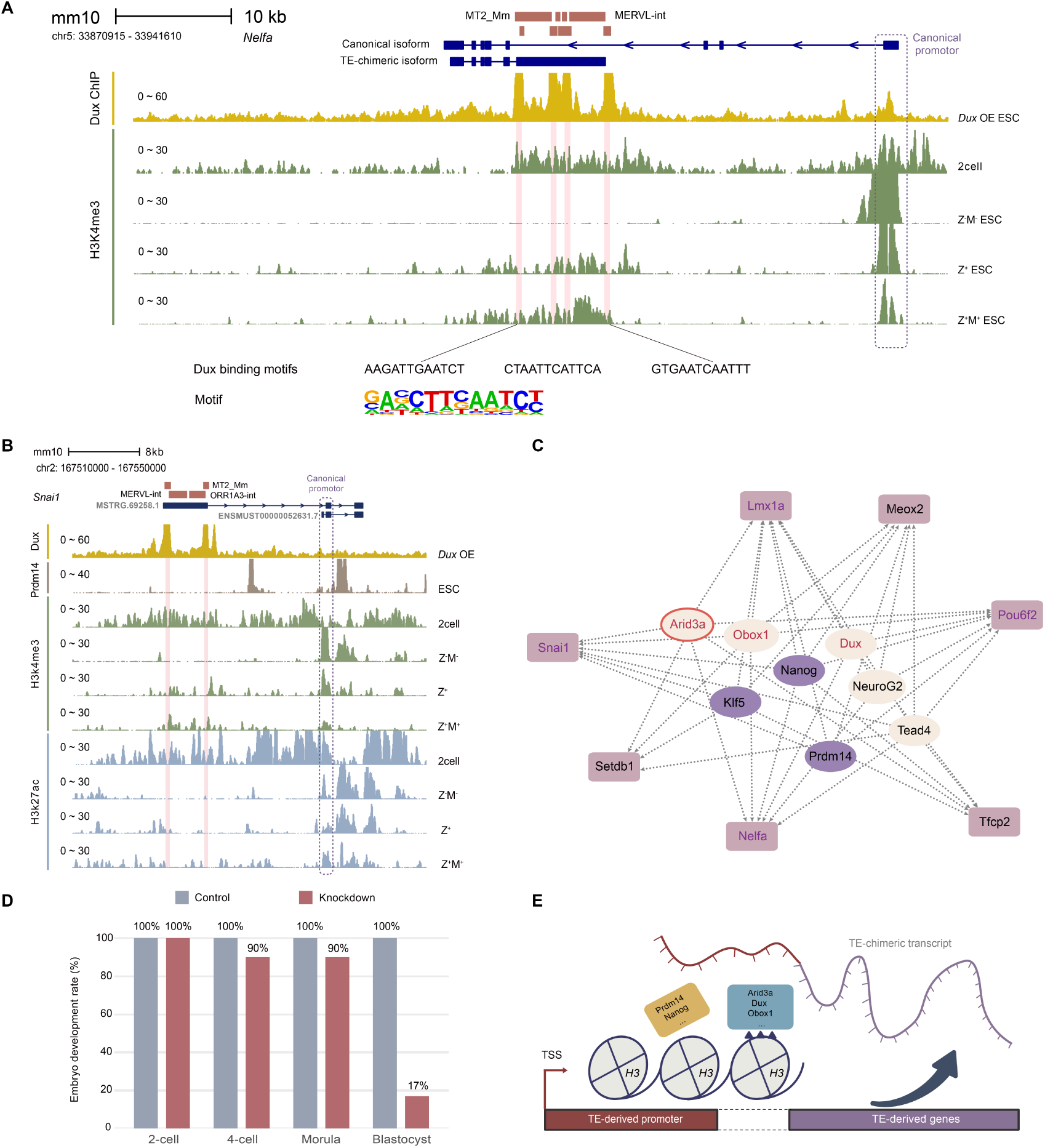
Regulatory Network Constructed by TEDDY Based on TE-chimeric Transcripts in 2C-like Transition. **A**, The genomic tracks of Dux ChIP-seq signals (yellow) and H3K4me3 signals (green) around *Nelfa* gene locus. These tracks illustrate the structure of the *Nelfa* canonical isoform and its truncated MT2-chimeric transcript isoform. The scale bar represents a 10 kb genomic span. The Dux binding sites, and corresponding motif are presented below. Z^-^M^-^: ESCs; Z^+^: *Zscan4*^+^ cells (intermediate cell state in the transition between ESCs and 2C-like cells); Z^+^M^+^: 2C-like cells. **B**, The track plot provides an overview of *Snai1*, delineating its TE-chimeric and canonical non-chimeric isoforms, as well associated epigenetic modification and TF occupancy around *Snai1* gene locus. The upstream TE acts as a potential promoter to drive *Snai1* chimeric isoform, which is extensively occupied by Dux. Z^-^M^-^: ESCs; Z^+^: *Zscan4*^+^ cells (intermediate cell state in the transition between ESCs and 2C-like cells); Z^+^M^+^: 2C-like cells. **C**, Regulatory network showing core TFs (central nodes) and driver TFs bearing TE-initiated transcripts (pink periphery nodes) in the 2C-like transition. Edges represent regulatory interactions between TFs and their targets, as predicted by TEDDY. Node colors denote potential function of TFs, with activators in yellow and repressors in purple. TF names shown in red indicate regulators supported by orthogonal evidence (Dux supported by ChIP-seq, Obox1 supported by prior functional studies, and Arid3a validated by perturbation), whereas purple gene labels denote experimentally validated TE-chimeric transcript targets. **D**, The embryo developmental rate of *Arid3a* KD embryos and WT embryos. The developmental rate was calculated as the ratio of embryos reach the indicated stage per number of zygotes. Two independent experiments were performed. **E**, Working model of TFs modulating TE-chimeric transcripts in 2C-like transition. Pluripotency-related TFs, including Nanog and Prdm14, inhibit TE-derived promoters for 2C-like transcripts and thus impede the transition from ESCs to 2C-like cells. Conversely, totipotency-related TFs including Dux, Obox1 and Arid3a act in an opposite manner to facilitate the generation of TE-chimeric transcripts and 2C-like transition. This model suggests that TE-derived promoters provide TF-binding substrates and are epigenetically regulated during 2C-like transition.

**Table 2.** Structure of the MT2B1-chimeric isoform of *Zfp352*.

| Transcript_id | gene_name | seqnames | start | end | width | strand | tx_exon_rank | TE_name | TE_class |
| --- | --- | --- | --- | --- | --- | --- | --- | --- | --- |
| ENSMUST00000107129.1 | Zfp352 | chr4 | 90218820 | 90218975 | 156 | + | 1 | MT2B1 | LTR |
| ENSMUST00000107129.1 | Zfp352 | chr4 | 90223583 | 90225702 | 2120 | + | 2 | none | none |

This regulatory logic extends to *Nelfa*, the gene encoding the upstream activator of *Dux*^42^. As noted above, we identified and validated an MERVL-initiated, N-terminally truncated *Nelfa* isoform (Fig. S5A). Strikingly, this TE-derived promoter shows strong Dux occupancy (Fig. 5A). H3K4me3 level (a marker of active promoters) increased in the intronic MERVL promoter but decreased in the canonical promoter region of *Nelfa* during the pluripotent-to-totipotent state transition (Fig. 5A). Concomitantly, Dux was preferentially bound to the intronic MERVL-derived promoter of *Nelfa*, rather than to its canonical promoter (Fig. 5A). These data collectively suggest a Dux-mediated switch from the canonical promoter to the TE-derived alternative promoter. TEDDY integrates transcriptomic and epigenomic data to outline the regulatory circuitry, nominating a potential Dux–*Nelfa* feedback loop in the establishment of totipotency.

To further explore the functional importance of TE-dependent transcripts, we turned to single-cell transcriptomics, analyzing published data to capture the transition from ESCs to 2C-like cells^52^. The UMAP analysis revealed a continuum of transcription states rather than distinct, well-separated clusters (Fig. S8A,B). We next performed pseudotime analysis to construct the reprogramming trajectory (Fig. S8C,D) and identified differentially expressed genes along the trajectory of 2C-like transition^53^. Based on these results, we identified TFs exhibiting significant expression changes during this process as potential regulators of 2C-like conversions (Fig. S8E). We categorized these TFs into two groups. TFs showing significantly higher expression levels in 2C-like cells relative to ESCs were defined as totipotency-associated TFs, whereas the TFs with opposite expression patterns were identified as pluripotency-associated TFs. Interestingly, we found that TE-derived promoters directly activate a number of totipotency-associated TFs (Table 3 and Fig. S8E). A representative example was *Snai1,* which generates an elongated chimeric isoform driven by the upstream MT2_Mm-derived promoter (Figs. S8E,F and S5A). Dux strongly occupied the MT2_Mm-derived promoter rather than the canonical one for *Snai1*, similar to the *Nelfa* example (Fig. 5B). Meanwhile, the canonical *Snai1* promoter exhibits a dramatic reduction in H3K4me3 level in 2C-like cells relative to ESCs (Fig. 5B). These findings from network inference and single-cell dynamics demonstrate that TE-chimeric transcripts are not peripheral outliers but constitute a core regulatory architecture. Of note, several well-known upstream regulators, such as *Nelfa* and *Zfp352*, are also driven by TE-derived promoters as mentioned above (Table 3 and Figs. 4H and 5A), reinforcing the role of TE-chimeric transcripts as key drivers of the totipotency network.

**Table 3.**
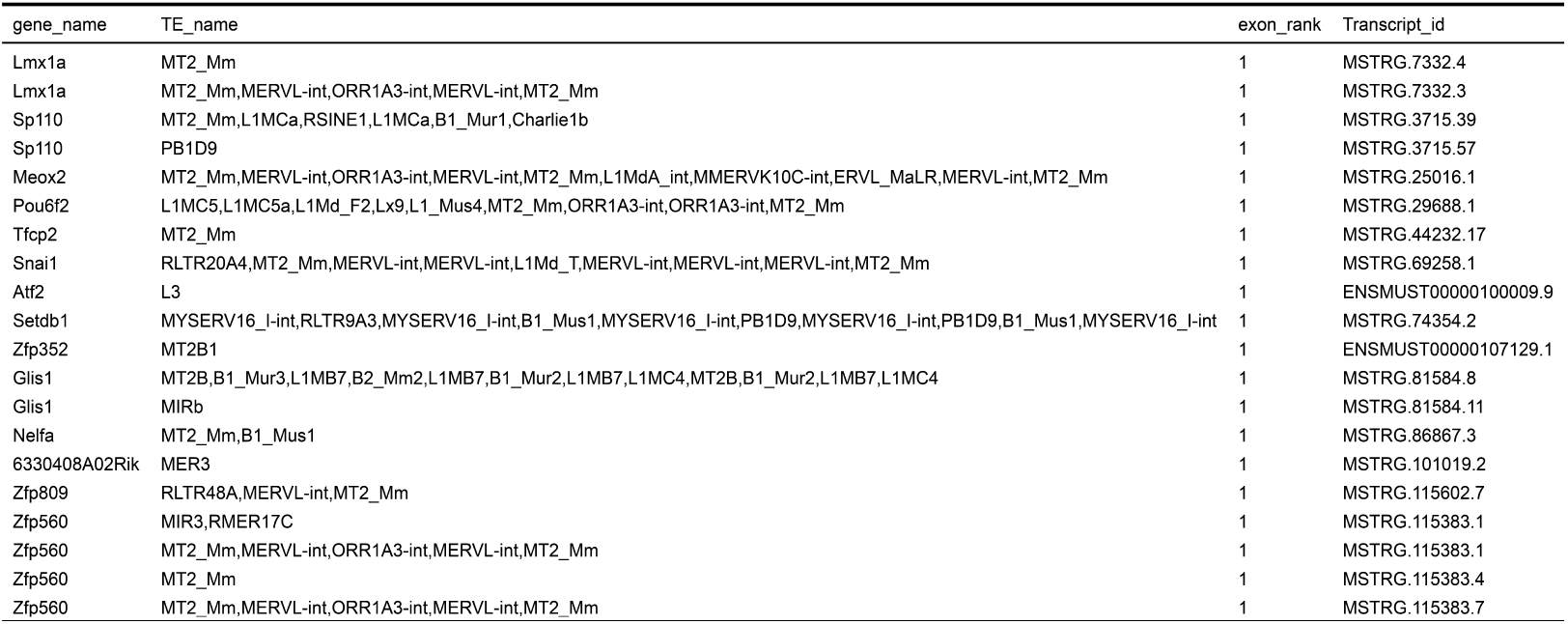
TE-chimeric isoforms of totipotency-associated transcription factors.

Motivated by the evidence for TE-promoter-mediated regulation, we further leveraged the isoform-resolved transcriptome incorporating TEDDY-detected TE-chimeric transcripts to apply the TEDDY Motif Search module (see Methods). Specifically, we tested whether candidate TF binding sites were present within the TE sequences associated with individual transcripts. These transcript-level TF–TE matches enabled the assembly of a comprehensive transcription factor regulatory network for the 2C-like entry transition (Fig. 5C). The resulting network systematically nominates regulatory relationships between TE-driven pioneer factors and the core totipotency- and pluripotency-associated TF network. This core set of regulators includes both well-known key factors, such as Obox1, and newly identified candidate regulators.

Several transcriptional targets of Obox1 predicted by TEDDY, such as *Hexb* and *Dppa4*, have been confirmed by recent functional studies^37,47,54,55^, supporting the biological plausibility of TEDDY’s motif-based nominations. These results move beyond the functional inferences from large-scale knockdown screens to pinpoint precise regulatory loci—defining both the TF binding sites (on TEs) and the specific target genes, thereby explaining how Obox1 activates ZGA genes (Table 4).

**Table 4.**
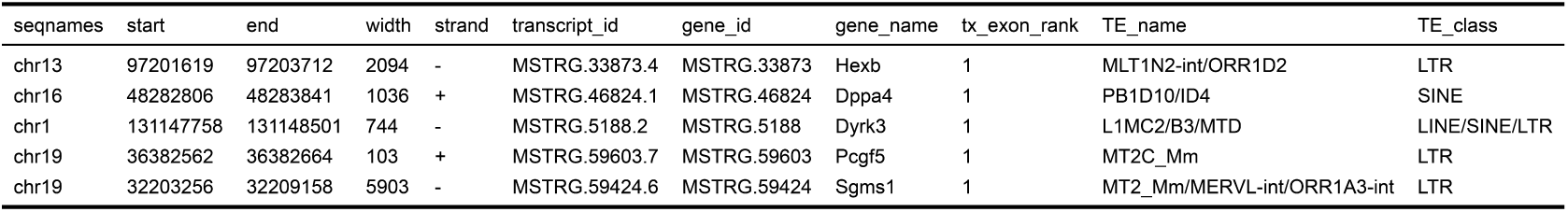
Experimentally validated Obox1-targeted TE-chimeric transcripts predicted by TEDDY.

To functionally validate the TE-chimeric regulatory architecture inferred by TEDDY, we focused on *Arid3a*, a top newly predicted regulator of the totipotency regulatory network. Overexpression of *Arid3a* facilitated the pluripotency-to-totipotency transition, as evidenced by upregulation of canonical hallmarks such as MERVL and *Zscan4* (Fig. S9B). Furthermore, it induced key transcription factors that activate the 2C-like state, such as *Dux* and *Obox1*. Notably, the TEDDY-identified TE-initiated isoform of *Zfp352* was strongly upregulated upon *Arid3a* overexpression, providing direct functional support for the regulatory logic inferred by TEDDY (Fig. S9C). These results indicate that *Arid3a* functions as a key regulator of totipotency establishment, acting through TE-mediated transcriptional regulation.

We next assessed the function of *Arid3a* in early development. We injected small interfering RNA (siRNA) targeting *Arid3a* into MII oocytes, followed by intracytoplasmic sperm injection (ICSI). A large fraction of *Arid3a-*KD embryos arrested at the morula stage (Figs. 5D and S9D), supporting an essential role for Arid3a during preimplantation development.

Previous studies have shown that *Nanog* inhibits 2C-like conversion and is degraded in 2C-like cells^56,57^. Together with the regulatory network for 2C-like transition constructed by TEDDY, these findings imply a potential antagonism between pluripotency-associated TFs and totipotency-associated TFs to modulate TE-associated promoters and the expression of associated 2C genes (Fig. 5C,E). Based on the isoform-resolved transcriptome, TEDDY enables locus-resolved inference of TF-target relationships by identifying candidate TF-binding sites within TE-derived regulatory elements and integrating them into refined TE-dependent regulatory hierarchies, revealing that core regulators such as *Arid3a* act through TE-derived promoters to activate downstream totipotency-associated TFs bearing TE-chimeric isoforms.

### Analysis of TE-chimeric transcription in human preimplantation embryos

Having established the role of TE-chimeric transcription in our 2C-like totipotency model system, we turned to staged human preimplantation embryos to assess the conservation of this regulatory mechanism *in vivo*. We therefore applied TEDDY to re-analyze RNA-seq data from human preimplantation development, including oocytes, 2-cell, 4-cell, 8-cell, and morula stages^58^.

TEDDY-resolved TE-chimeric transcripts were visualized as a heatmap. This overview reveals a broad and dynamic landscape of TE-driven transcription in early human development, with a distinct stage-specific pattern similar to, but not identical with, that observed in mouse early embryos (Figs. 6A and 3A). Early human samples, notably oocytes and pronuclear-stage embryos, are enriched for distinct TE-chimeric isoforms (Fig. 6A). As embryos progress toward embryonic genome activation, new TE-chimeric clusters emerge, including a small cohort of 4-cell specific transcripts and a broader pre-8-cell shared wave that fades before the 8-cell stage. The most pronounced transition occurs at the 8-cell stage, followed by a rapid decline from the morula stage onward, coinciding with major ZGA (Fig. 6A)^59^. Another set of TE-chimeric transcripts shared between 8-cell and morula remains highly expressed, marking the post-ZGA regulatory state (Fig. 6A).

**Fig. 6.**
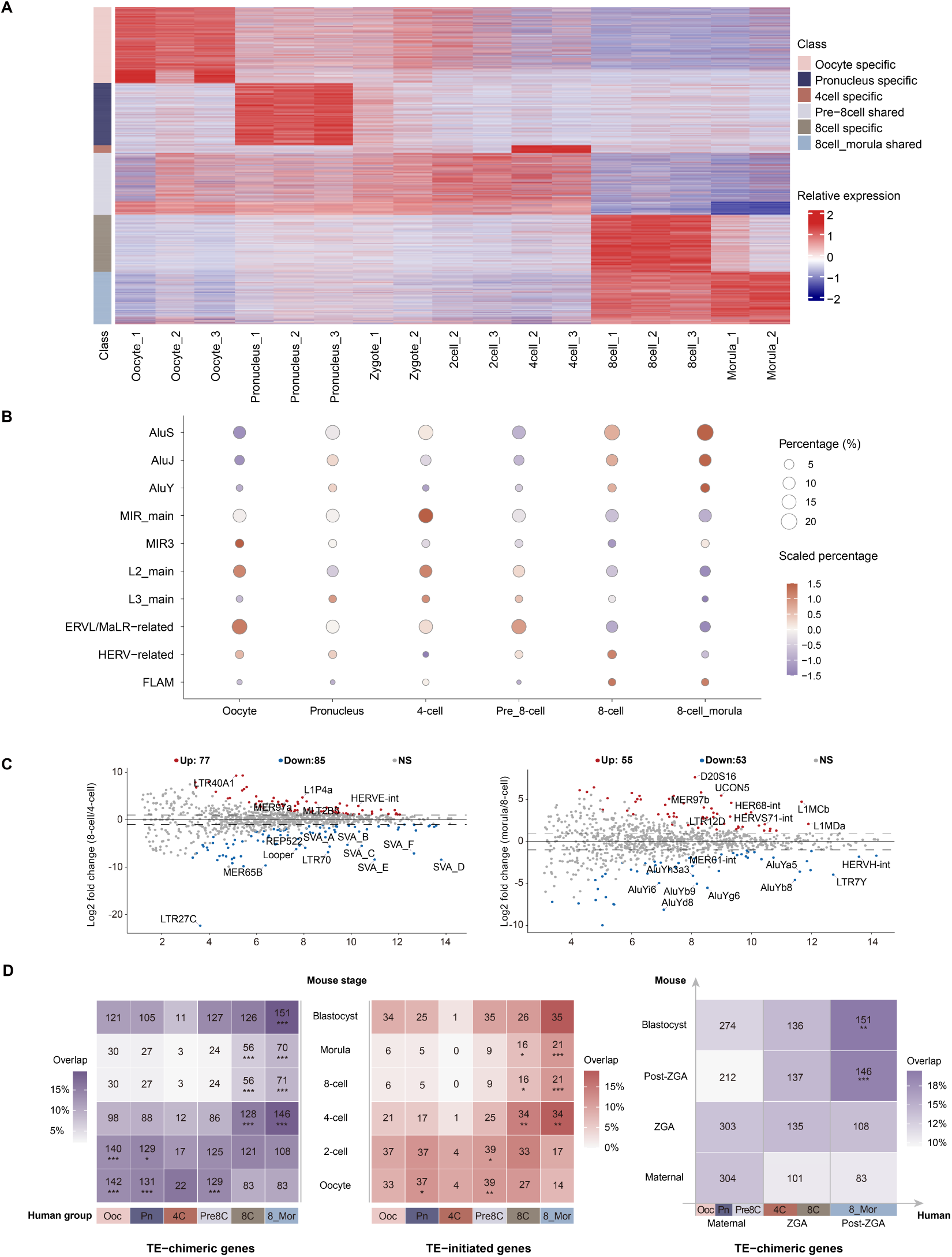
Characterization of TE-chimeric Transcripts in Human Preimplantation Development. **A**, Heatmap showing row Z-score normalized expression of TE-chimeric transcripts across seven embryonic stages, from oocyte to morula. Unsupervised clustering resolves six classes with distinct stage-specific patterns, particularly at the oocyte, pronucleus, 8-cell and morula stages. Each row represents the Z score of log2-transformed FPKM values. **B**, Dot plot showing the proportions of prominent TE categories associated with TE-chimeric transcripts in the developmental expression groups identified in Fig. 6A. Dot size indicates the absolute proportion of each TE category within each group, whereas color represents the row-scaled proportion, calculated across developmental groups for each TE category. TE subfamilies were collapsed into broader categories, including Alu, MIR, LINE, ERVL/MaLR-related, and HERV-related groups. **C**, Differential abundance of TE subfamilies across human pre-implantation transitions. MA plots show log2 fold change (y-axis) versus log2 mean abundance (x-axis) for 8-cell vs 4-cell (left) and morula vs 8-cell (right). Points are colored by significance (red, up; blue, down; grey, not significant). **D**, Heatmaps comparing human TE-associated expression groups with mouse developmental stage-associated TE-chimeric or TE-initiated gene sets via mouse-human orthologs. (Left, Middle) Overlap of TE-chimeric and TE-initiated genes, respectively. (Right) Aggregated developmental-wave analysis of TE-chimeric genes. Mouse stages are grouped around the 2-cell ZGA transition (maternal, ZGA, post-ZGA, and blastocyst waves), and human groups are collapsed into broader waves (see Methods). The bottom color bars indicate human expression groups from Fig. 6A: Ooc, Oocyte specific; Pn, Pronucleus specific; 4C, 4cell specific; Pre8C, pre-8cell shared; 8C, 8cell specific; 8_Mor, 8cell_morula shared. Tile colors represent the overlap ratio (the proportion of human genes per group/wave whose mouse orthologs show TE-associated activation in the corresponding mouse stage/wave), with numbers indicating the overlapping ortholog count. Asterisks denote preferential overlaps based on mouse-stage-anchored Pearson chi-square residual analysis (Left, Middle) or significant hypergeometric enrichment after Benjamini–Hochberg correction (Right).

We next used TEDDY to analyze the structural details and TE family/subfamily origins of TE-chimeric transcripts at each developmental stage (Fig. 6B). The results show a heterogeneous but stable background dominated by Alu subfamilies, MIR elements and L2s throughout preimplantation. Compared to mouse early embryos, human embryos use a different and more complex repertoire of TEs in TE-chimeric events (Figs. 3B and 6B,C), with a much greater contribution from SINE and LINE elements. This suggests that the stage-specific activation of TE-chimeric transcription is shared between mouse and human, whereas the specific TE families recruited into this process are species-specific. These differences may partly stem from the distinct genomic composition and evolutionary history of TE families in the two species, including the high prevalence of Alu and SINE elements in the human genome and the prominent contribution of LTR/ERV elements in mouse^60^. We further asked whether the TE subfamilies contributing to 8-cell specific TE-chimeric transcripts were themselves preferentially enriched at the 8-cell stage. Among ∼1,700 8-cell specific TE-chimeric events, only 2.02% originated from subfamilies enriched in 8-cell embryos over both 4-cell and morula stages; ∼28% were associated with subfamilies enriched over only one adjacent stage; and the remaining 69.8% involved subfamilies whose bulk abundance at the 8-cell stage was not elevated relative to either of the other two stages (Supplementary Table 6). Conversely, for TE subfamilies highlighted as 8-cell enriched in the abundance analysis (Fig. 6C), many TE copies were located within annotated gene bodies and within genes expressed at the 8-cell stage, but only a subset of these loci produced detectable TE-chimeric transcripts, and an even smaller subset produced TE-chimeric transcripts expressed at the 8-cell stage. Thus, TE abundance, genomic occurrence and host-gene expression are not sufficient to account for TE-chimeric transcript formation. Together, these results indicate that 8-cell-specific TE-chimeric transcription is temporally regulated rather than a passive consequence of increased bulk TE subfamily abundance.

We next compared TEDDY-identified TE-chimeric and TE-initiated events between mouse and human preimplantation embryos at the orthologous gene level to examine their cross-species recurrence (Fig. 6D). For each human expression group defined above, we quantified its overlap with mouse stage-associated TE-chimeric or TE-initiated gene sets across all human expression group-mouse stage combinations, without assuming one-to-one developmental stage correspondence. Stage-associated preferential overlaps were assessed statistically as described in Methods. Overall, TE-chimeric genes and the narrower TE-initiated subset showed broadly similar cross-species overlap patterns. The overlap did not follow a strict one-to-one correspondence between mouse developmental stages and human expression groups. Instead, human early expression groups showed preferential overlap with mouse oocyte and 2-cell gene sets, whereas human 8-cell- and morula-associated groups showed stronger overlap with later mouse stages. Several earliest human expression groups preferentially overlapped with mouse oocyte gene sets, prompting us to explore representative ortholog-level examples of potential old or recurrent TE-associated activation during the early maternal phase of development (Supplementary Table 7). Only a few candidates used the same ancient TE subfamily as the TE-derived promoter, including MIR, MLT1D, MER5A, L2b and L2c^61^. Other candidates instead involved different TE sources between mouse and human, such as mouse MTA_Mm versus human LTR12F, THE1 and other elements.

Given the more distributed timing and TE-source usage of human TE-chimeric transcription compared with the sharp LTR/ERV-dominated ZGA in mouse, we further collapsed the human expression groups and mouse developmental stages into broader maternal, ZGA, and post-ZGA waves. Human post-ZGA-associated TE-chimeric genes showed the strongest enrichment with mouse post-ZGA and blastocyst-stage gene sets (see Methods), indicating a degree of orthologous gene-level conservation after ZGA. In contrast, ZGA-associated TE-chimeric transcription appeared more species-specific, mainly drawing on lineage-specific TE repertoires, including Alu/HERV-related elements in human (Fig. 6B) and LTR/ERVL-related elements in mouse (Fig. 3). Overall, human TE-chimeric transcription is deployed more gradually across preimplantation development, whereas mouse TE-chimeric transcription is concentrated around a sharp ZGA-associated burst.

### TE exonization shapes isoform diversity and associated clinical outcome in hepatocellular carcinoma

Transposable elements have attracted growing interest in cancer research^30,62^. Increasing evidence indicates that TEs contribute to tumor-associated transcriptional and genomic alterations via diverse mechanisms. They may promote genomic instability through insertional events and rearrangements, and can be co-opted as cis-regulatory sequences to rewire transcription^30,62^. Importantly, recent work has shown that activation of TE-derived promoters can drive oncogene overexpression, activate tumor-suppressive pathways^63^ and produce TE–gene chimeric proteins, with potential implications for tumor biology and immunogenicity. These developments motivate systematic, transcript-level characterization of TE-driven transcriptional events in tumors.

Hepatocellular carcinoma (HCC) arises from diverse etiologies, such as HBV/HCV infection, alcohol and aflatoxin exposure^64^. Moreover, HCC typically shows abundant alternative splicing and isoform diversity^65^ alongside substantial TE burden, which poses both a challenge and a need for accurate isoform reconstruction avoid ambiguous assignment of TEs to adjacent genes. For example, a recent study showed that liver TEs recruit KDM1A to repress neighboring *HNF4A* and ultimately promote HCC tumor growth^66^. However, that analysis primarily relied on proximity-based matching, limiting its ability to clarify potential TE–gene relationships and exon-chain structure.

Thus, HCC provides a particularly well-suited testbed for methods targeting TE-chimeric events. We first applied TEDDY to a cohort of HCC samples to detect, quantify and integrate TE-chimeric transcripts. Specifically, we analyzed RNA-seq data from five HCC patients^67^, three with primary tumors and two with relapsed disease, representing both sexes and a range of ages. Each patient had matched adjacent-normal and tumor tissue. To illustrate our multi-sample capabilities, we summarized the presence and absence of TE-chimeric transcripts across the ten libraries using the pipeline’s integrated visualization (Fig. 7A,B). The UpSet analysis indicates that most detected TE-chimeric transcripts are reproducibly observed across multiple samples, revealing a core shared set of events. Smaller sample- and patient-specific intersections are also apparent. Several transcripts are relatively enriched in relapse samples, a pattern that may point to disease-associated transcriptional shifts. TEDDY further highlights TE-initiated events (Fig. 7A) and the contribution of LINE-derived sequences (Fig. 7B) among TE-associated isoforms, facilitating rapid assessment of the overall distribution and cohort-level patterns of TE-chimeric transcription. We further extended the structural analysis to two independent pairwise tumor-normal HCC datasets^68^ and observed a similar overall structural distribution. Across all three HCC cohorts, comprising 75 patients and 150 samples in total, approximately one quarter of L1-chimeric transcripts were classified as LINE1-initiated, whereas the majority represented LINE1 internally embedded in the mRNA sequence, reflecting TEDDY’s capacity for large-scale cohort-level transcript structure annotation.

**Fig. 7.**
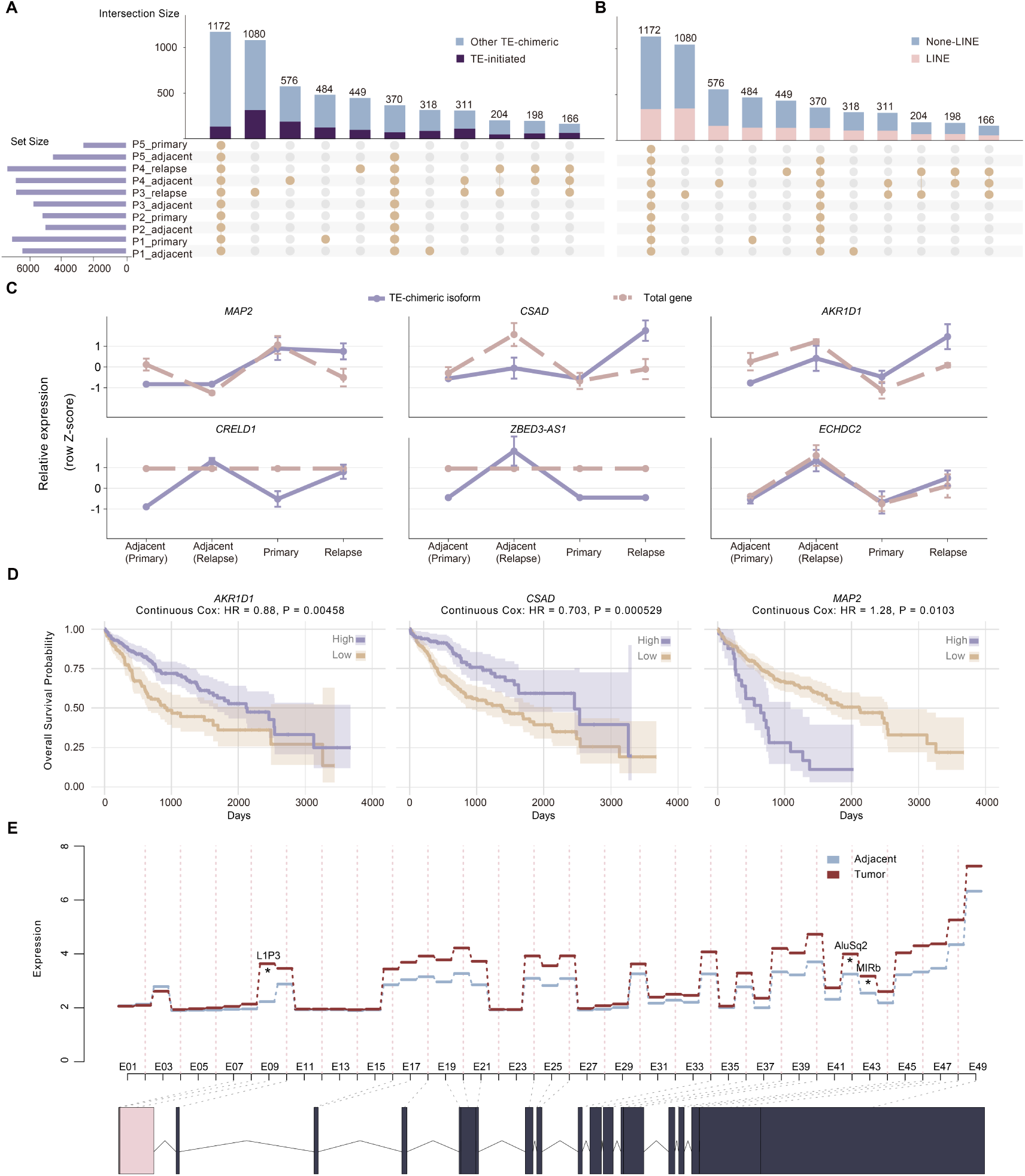
Landscape and Clinical Relevance of TE-chimeric Events in HCC. **A**, UpSet plot showing shared and unique TE-initiated transcripts across ten RNA-seq libraries from five HCC patients with matched tumor and adjacent normal samples. Transcripts identified as TE-initiated are highlighted in purple. Vertical bars indicate the number of transcripts shared by the sample combination denoted by the connected dots below each bar. The horizontal bar plot on the left shows the total number of TE-chimeric transcripts detected in each individual sample. **B**, Distribution of LINE1-derived transcripts. The UpSet plot follows the same sample configuration and set sizes as panel A, with transcripts harboring LINE1-derived sequences highlighted in pink. **C**, Representative TE-chimeric isoform and host-gene expression patterns in the four-condition HCC cohort. Line plots show the mean relative expression of selected TE-chimeric isoforms and their corresponding total host genes across adjacent tissues from primary HCC patients, adjacent tissues from relapse HCC patients, primary tumors and relapse tumors. TE-associated isoform expression and VST-transformed total host-gene expression were separately standardized within each isoform or gene to Z-scores for visualization, and error bars indicate SEM across biological replicates. **D**, Kaplan-Meier curves for *MAP2*, *CSAD* and *AKR1D1* in TCGA-LIHC. Survival association was evaluated using continuous Cox regression with log_2_(FPKM + 1) transformed gene expression. HR corresponds to a one-unit increase in this transformed expression, and Cox *P* values are shown above each plot. Patients were stratified into high- and low-expression groups using an optimal expression cutpoint for visualization. **E**, Step plot showing exon usage and normalized RNA-seq read counts of individual flattened exon bins of *MAP2* in 65 tumor-adjacent pairs from HCC patients. Adjacent and tumor samples are shown in blue and red, respectively. The x-axis displays flattened exon bins across the *MAP2* gene range. TE-associated bins identified by TEDDY as significantly differentially used between tumor and adjacent samples are marked with asterisks. The bottom diagram illustrates the structure of the representative L1P3-initiated *MAP2* isoform, with dotted lines linking its included exon bins.

Applying TEDDY’s chimeric-driven test to the initial HCC dataset, we identified a set of TE-chimeric isoforms that exhibit reproducible, condition-dependent patterns. A subset of these candidates is supported by prior literature implicating them in liver cancer biology. This set includes key players such as *AKR1D1*, a central regulator of bile acid metabolism recently shown to suppress liver cancer progression by bridging metabolic dysregulation and NK cell immunity^69^. We also identified *CSAD*, whose alternative splicing is regulated by the splicing factor *RBM17* to promote tumorigenesis^70^. The relapse-adjacent enriched TE-chimeric isoform of *ZBED3-AS1* is also notable in light of previous reports that *ZBED3* overexpression can reactivate the Wnt/β-catenin signaling pathway to promote tumorigenesis^71^.

However, host-gene expression did not uniformly follow these isoform-level shifts, indicating that differential usage of TE-chimeric bins does not necessarily translate into proportional changes in total gene-level abundance. We therefore further evaluated the contribution of TE-chimeric isoform usage to host-gene expression patterns in the HCC data. Specifically, we defined a post-hoc isoform-gene concordance score integrating TE-chimeric isoform expression, TE isoform ratio and total host-gene expression (see Methods). The concordance score captured the distinct patterns observed in Fig. 7C. *MAP2*, *CSAD* and *AKR1D1* showed high concordance scores, reflecting TE-chimeric isoform upregulation, increased TE-associated isoform ratios, and their potential contribution to concordant changes in total host-gene abundance. By contrast, *CRELD1* and *ZBED3-AS1* showed TE-chimeric isoform-level changes without robust gene-level perturbation. *ECHDC2* represented a context-dependent TE-exonized event, with an increased TE-associated isoform ratio in relapse-adjacent samples and only modest concordant changes in host-gene abundance, rather than a strong gene-level drive-like pattern.

We next applied TEDDY to two independent pairwise tumor-normal HCC cohorts, performing TE-chimeric bin differential-usage analysis and calculating an isoform-gene concordance score. This identified 26 genes that exhibited differential usage of TE-chimeric bins together with recurrent patterns of TE-driven expression variation across three independent HCC cohorts. We therefore asked whether genes with such recurrent TE-linked expression variation were associated with patient survival. Taking HCC TE-chimeric genes included in the TCGA-LIHC Cox analysis as the background, these three-cohort-supported genes were significantly enriched among survival-associated genes (Fisher’s exact test, odds ratio = 3.18, *P* = 0.0053). Specifically, continuous Cox regression in TCGA-LIHC primary tumor samples showed that higher *MAP2* expression was associated with poorer overall survival, whereas higher *CSAD* and *AKR1D1* expression was associated with better overall survival (Fig. 7D). Notably, *MAP2* harbored a representative novel L1P3-initiated TE-chimeric isoform, independently identified across all three HCC cohorts. This isoform was also supported by three independent TE-chimeric bin differential-usage analyses and exhibited consistent tumor-associated induction (Fig. 7E).

## Discussion

The activity of transposable elements (TEs) is increasingly observed across diverse biological contexts. Notably, the functional incorporation of TEs into host genes significantly diversifies the transcriptome, yet the fundamental principles of TE-driven regulation remain largely elusive, highlighting the need to systematically profile these events. Thus, we developed TEDDY, an end-to-end computational solution for the detection, reconstruction, quantification, and downstream analysis of TE-chimeric isoforms. For a method designed to address an underlying biological challenge, benchmarking should not be restricted to purely in silico metrics or algorithm-level comparisons. We therefore adopted a multi-level validation strategy, establishing baseline detection accuracy through controlled in silico simulations, confirming technical robustness across diverse sequencing platforms, and demonstrating structural resolution and cross-species generalizability via locus-by-locus validation in *Drosophila*. Ultimately, and most importantly, we evaluated whether TEDDY could be applied in authentic biological contexts to generate and validate biological discoveries.

Enabled by TEDDY’s capacity for large-scale, parallel sample analysis, the resulting atlas of TE-chimeric transcripts revealed a widespread temporal switching during the preimplantation development of both mouse and human embryos, involving diverse TE families that exhibit distinct, stage-specific and species-specific expression patterns. Considering their potentially critical roles in early embryogenesis, we further focused on 2-cell embryos and 2C-like cells to model totipotency, confirming that the *in vitro* model recapitulates the extensive TE-chimeric transcriptional landscape of its *in vivo* counterpart. Within this system, TEDDY systematically uncovered that core totipotency factors (e.g., *Nelfa*, *Snai1*) undergo TE-derived promoter switching to generate previously uncharacterized isoforms. Crucially, independent validation via RT-PCR, Sanger sequencing and third-generation long-read sequencing confirmed both TEDDY’s computational accuracy and the reliability of these biological findings, substantially refining the widely studied totipotency network.

Building on the dual-level structure and quantification of TE-chimeric transcripts, specifically capturing differential TE-chimeric bin usage and broader isoform switching, we integrated multi-omics data to dissect how well-known pioneer factors, such as *Dux* and *Obox1*, orchestrate the totipotency network through TE-chimeric-transcript-mediated regulation. While traditional approaches often rely on large-scale perturbations or imprecise genomic proximity to infer correlative associations, TE-chimeric sequences provide a physical clue to explicitly construct locus-resolution TF-TE-target gene relationships. By tracing potential upstream TFs and their exact targets, we constructed a putative totipotency network and nominated several novel core regulators, including *Arid3a*. Its overexpression potently activated totipotency-associated markers and TFs, while its embryonic knockdown resulted in developmental arrest. This validation not only establishes TE-chimeric regulation as a core mechanism essential for early embryonic development, but also demonstrates TEDDY as an efficient framework for decoding the pervasive role of TE-chimeric transcription.

Developmental programs are frequently reactivated in cancer and can be co-opted to support proliferation, plasticity and immune evasion^72^. Three independent hepatocellular carcinoma (HCC) cohorts demonstrate TEDDY’s scalability for parallel analysis of large-scale, heterogeneous transcriptomic datasets. Within these cohorts, we identified recurrent and clinically informative TE-chimeric transcripts, providing attractive targets for further mechanistic follow-up.

As sequencing technologies advance across generations, isoform- and TE-associated studies still represent a vast, underexplored frontier. As a discovery framework, TEDDY prioritizes candidate TE-dependent isoforms and regulatory relationships, whereas the causal roles and molecular mechanisms of individual events require locus-specific perturbation and functional validation in the relevant biological context. We anticipate that TEDDY will serve as a broadly applicable framework to integrate multi-platform data and comprehensively explore TE-dependent transcription across development, evolution, and disease.

## METHODS

### Transcriptome reconstruction

The functional interpretation of TE-dependent isoforms requires full host-gene context, making genome-coordinate exon-chain reconstruction essential. Thus, TEDDY adopts a reference-guided transcriptome reconstruction approach with local *de novo* isoform discovery (see Supplementary Note 1). To achieve this, the alternative splicing graph model of StringTie serves as a natural and suitable foundation^73,74^. TEDDY integrates the StringTie framework as its core reconstruction engine, providing an end-to-end analysis workflow within the R environment (Fig. 1A). The pipeline accepts splice-aware alignments from short-read (e.g., Illumina) and third-generation long-read (e.g., ONT, PacBio) platforms, together with a reference annotation, as initial inputs. This initial reconstruction step generates isoform-resolved transcript models for each sample, capturing genomic coordinates and exon-chain structures, including the recovery of novel unannotated isoforms. Because transcript reconstruction occurs independently across samples, the resulting per-sample annotations possess sample-specific identifiers and inconsistent structural boundaries, precluding direct cross-sample comparison. Alternative splicing graphs (ASGs), originally used to assemble transcript structures from sequencing reads, are also well suited for cross-sample isoform harmonization, because transcript models from different samples often share the same intron chain but differ slightly at transcript boundaries. TEDDY therefore invokes the StringTie graph framework to collapse such structurally concordant isoforms into a unified non-redundant reference. The merged transcript set retains genomic coordinates but lacks standardized gene context and transcript-class annotation. TEDDY then annotates the merged assembly against the reference annotation using GffCompare^75^. This step assigns gene correspondence and transcript class codes, while further collapsing redundant models that share identical intron chains but differ only slightly at their boundaries, thereby yielding a more concise set of exon-chain structures and an interpretable unified transcript reference.

### Identification and quantification of TE-chimeric transcripts

TEDDY annotates the unified reconstructed transcriptome assembly GTF file via the transposon annotation obtained from RepeatMasker^76^. Accurate exonic structure provides the fundamental analytical abstraction for alternative splicing and TE-chimeric transcript analysis^77^, because the relevant variation is manifested primarily through exon composition and transcript boundaries rather than base-by-base sequence differences^78^. TEDDY addresses this by organizing reconstructed transcript structures into ranked exons and, for bin-level quantification, further flattening the exon structures of all isoforms within each gene locus into disjoint exonic segments used as counting bins^79,80^, as illustrated in Fig. 1A. The transposon-derived exons or bins are then annotated using the transposon annotation mentioned above. Therefore, we can determine the specific TE exonization events, along with the corresponding transposons and the sequence of the fusion transcripts. TEDDY provides both transcript-level and the counting-bins-level quantification and annotation among multiple samples, ensuring structured data storage in two SummarizedExperiment objects for subsequent analyses. For TE-bin quantification, TEDDY summarizes reads from splice-aware BAM alignments using default counting settings. Based on the previously generated unified GTF file and the positional coordinates of exons in the metadata of Summarized Experiment object, users can extract the reference sequence of the transcript. This serves as an important resource for conducting subsequent investigations into the transcript sequence and exploring its potential implications on protein functionality.

### Detection of differential usage of chimeric exons

Differential usage of TE-chimeric exons serves as an indicator of whether a TE-containing isoform becomes dominant under specific conditions, providing insight into its regulation and functional implications.

TEDDY adapted the statistical method of DEXSeq^80^ and developed an algorithm to examine the impact of fusion transposable elements on gene expression. We define N_mnl_ as the count of reads falling into the counting bin l of gene m in sample n. Each count N_mnl_ is treated as an outcome of a random variable, assumed to follow a negative binomial distribution *N_mnl_* ∼ NegBin(μ*_mnl_*, *θ*), where *μ_mnl_* is the mean and *θ* is the dispersion parameter. Our null hypothesis posits that the relative expression of the TE-chimeric bins does not fluctuate under different conditions, thus not contributing to significant variations in gene expression. For each gene, we fit a generalized linear model (GLM) to the count data N using the variables S for the sample effect, C indicating whether the bin is TE-chimeric or non-chimeric, and X denoting the experimental conditions.

For each bin l that incorporates a TE sequence, we fit a model to assess how the presence of TE sequences influences the read count distribution across different conditions. This model setup allows us to observe how the fraction of reads overlapping this TE-chimeric bin varies in proportion to the reads overlapping the gene under specific conditions as in Equation (1):

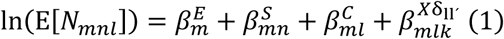

Here, 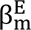 represents the baseline expression across all bins for gene m, 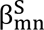 estimates the logarithm of the fold change to mitigate sample effects across all bins for gene m, and 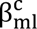 adjusts for the expression impact introduced by TE sequence incorporation, differentiating between TE-chimeric or non-chimeric bins for gene m. The term 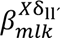, where δ_lĺ_ is 1 if l = ĺ and 0 otherwise, specifically tests the interaction for this TE-chimeric bin under varying conditions.

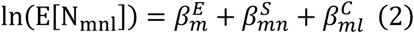

This null model in Equation (2) excludes the interaction term and is used to determine whether the main effects explained by the three terms are sufficient to account for the expression variability across different conditions, thereby revealing the significance of the interaction effects. TEDDY then calculates the difference in deviances between the two fits and conducts a chi-square test to obtain a p-value, assessing whether the null hypothesis is supported. If the null hypothesis is rejected, the model with interaction terms provides a better explanation, indicating significant differences in the TE-exon under different conditions.

To implement this analysis, TEDDY proceeds based on the previously mentioned bin-based expression data and TE-chimeric annotation. Initially, TEDDY calculates size factors using the median ratio method to adjust for differences in library size or sequencing depth across samples^81^. To estimate the dispersion parameters essential for accurate differential expression analysis, we utilize several functions from the DESeq2 package in TEDDY. The process includes Initial Estimation, Model-Based Refinement, and Maximum A Posteriori (MAP) Estimation^82^. With normalized data and estimated dispersion parameters in place, TEDDY fits both the null model and the full model to the data using the function ChimericDrivenTest, and results can be extracted using extractTest. TEDDY also encompasses the function calculateFoldchange to extract various coefficients of the selected gene from the GLM fit. In this context, the GLM fit refers to the multi-factor fitting of negative-binomially distributed read counts to isolate the contribution of TE-chimeric bin usage from the baseline gene transcription. This functionality enables users to gain valuable perspectives into the perturbation patterns of the target gene’s expression, as influenced by different conditions and the presence of TE components.

TE-associated bins comprised both TE-initiated and internal TE-exonized segments defined by TEDDY’s chimeric annotation. The design matrix included the relevant experimental condition (e.g., cell state or disease group) and, where applicable, library or batch effects as fixed factors. For each gene, TE effects were tested by comparing a full negative-binomial GLM (log link; including TE-bin term[s]) against a null design (without TE terms) using a likelihood-ratio test (nbinomLRT). *P*-values were adjusted using the Benjamini–Hochberg procedure. Features with all-zero counts or zero counts in the corresponding “others” component were excluded because no valid test statistic could be computed. By default, bins passing a user-configurable BH–adjusted *P*-value threshold were reported as TE-driven usage events (in this study, BH–adjusted *P* < 0.01 unless stated otherwise). These events denote condition-dependent shifts in the inclusion of TE-associated sequences, consistent with underlying changes in TE-chimeric isoform usage.

### Quantitative visualization of transcript structure

To support interpretation of TE-chimeric isoforms and their condition-dependent usage changes, TEDDY provides plotting functions that operate directly on transcript-level and bin-level SummarizedExperiment objects. Two complementary views are generated. (i) Isoform view: a compact transcript model showing the exon–intron structure for a selected TE-chimeric transcript, with TE-overlapping exonic segments highlighted according to the chimeric annotation. (ii) Gene-locus view: an exon/bin– resolved profile summarizing normalized bin-level counts for a gene across conditions, with bins identified as significant by the differential usage test emphasized. When needed, the locus view can be anchored to a specific transcript by rendering the corresponding isoform model beneath the gene track, enabling direct alignment between statistical evidence at the bin level and transcript structural context. Together, these plots provide a consistent bridge from bin-level inference to locus-level structural interpretation.

### Regulatory network inference based on TE-chimeric transcripts

Previous regulatory analyses, including HOMER-based analyses, have commonly relied on motif enrichment across pooled genomic regions derived from large-scale omics datasets and inference based on genomic proximity. The TE-chimeric junction serves as a structural bridge linking a specific upstream TE-derived cis-regulatory sequence to its host gene. Reconstruction of TE-chimeric transcripts therefore enables regulatory inference at transcript and locus resolution. TEDDY includes a motif-scanning module for transcript-specific regulatory inference and retrieves the full sequence of the TE associated with each TE-chimeric transcript. Candidate binding sites for transcription factors (TFs) of interest are then systematically scanned across these sequences using position weight matrix matching. This procedure generates transcript-resolved TF–TE matches linking TFs, TE-derived regulatory elements, and downstream host genes, yielding sequence-level support for putative regulatory networks and locus-specific motif-hit information for downstream analysis and experimental validation.

### Software implementation

TEDDY is provided as an R package and integrates optimized C++ routines via Rcpp for computationally intensive steps. Sample-level parallelization is supported to enable efficient processing of multi-sample datasets. TEDDY orchestrates external tools for specific pipeline stages and also implements built-in modules for TE-exonization detection, TE annotation, isoform-aware quantification and isoform-switching analysis, visualization functions, and reconstruction of putative TE–gene regulatory networks. It also provides wrapper functions to standardize inputs and outputs. For code availability and the exact software release used in this study, see the Code availability statement.

### In silico Benchmark design

#### Simulation Setup

To construct ground-truth annotations, we derived simulated GTFs from the mouse GENCODE M7 reference annotation. We first identified exons with promoter potential and treated these as putative transcriptional start sites. For each selected start exon, we concatenated a small number of randomly selected downstream exons from the same gene in a strand-consistent manner to generate biologically plausible transcripts. The simulated GTFs were then annotated to capture transcript composition and classify chimeric events. We next generated *in silico* RNA-seq datasets using RSEM. We sampled transcript abundance profiles from real embryonic RNA-seq data to preserve empirical abundance distributions and fragment-length characteristics. We generated paired-end reads at five nominal coverage levels (5×, 10×, 25×, 50× and 100×) and applied ∼10% stochastic perturbation with RSEM’s sampling and sequencing-error model to emulate read-level variability. Starting from the simulated ground-truth GTF, we randomly selected 90% of transcript models to form a partial reference, mimicking incomplete real-world annotations.

#### Benchmark evaluation and scoring

All tools were executed according to their recommended guidelines, with tool-specific settings and output filters summarized in Supplementary Table 1. To enable a fair comparison across tools with native parameter constraints and output formats, benchmark outputs were harmonized at a common host-gene level but compatible truth sets were matched to tool scope (Supplementary Table 1). Specifically, TEProf2 and LIONS were evaluated against simulated TE-initiated genes, whereas FREDY was evaluated against genes satisfying its native strand-unaware ≥ 50% TE-overlap criterion. TP, FP, and FN were derived by comparing predicted host-gene sets with the corresponding truth sets. Precision(*P*), recall(*R*), and the *F*_1_ score were defined as P = TP/(TP + FP), *R* = TP/(TP + FN), and *F*_1_ = 2*PR*/ (*P* + *R*), respectively.

#### Read-support in simulated data

To assess whether simulated TE-chimeric breakpoint structures were represented by aligned reads, we evaluated read-level support from aligned BAM files generated from the simulated RNA-seq reads. Because TE-overlap segments differ in length and local topology, we categorized them based on the length of the TE sequence incorporated into the mRNA and applied both continuous-alignment and anchor-pair support criteria accordingly. TE-overlap segments shorter than 125 bp and embedded within an exon were classified as short TE full-span events, because the TE-derived segment and adjacent host sequence could in principle be covered by individual reads. For these events, continuous-alignment support required an individual read to align continuously across the full TE-overlap segment together with adjacent host sequence. Those of at least 125 bp were classified as long TE boundary events, for which continuous-alignment support required an individual read to span the junction region. For both classes, anchor-pair support required shared read names between TE-side and host-side anchor regions, corresponding to paired-end read linkage, with a minimum alignment overlap of 15 bp for each anchor. Support rates for expressed truth loci were summarized across sequencing depths and, to assess the expected effect of transcript abundance, further stratified by pre-assigned TPM values within the 100× dataset.

### Bulk RNA-seq preprocessing

Raw reads from all platforms were processed using TEDDY’s built-in WDL workflow to produce splice-aware, sorted BAM files for reconstruction. Illumina RNA-seq reads were aligned with STAR v2.7.9a using the default multi-mapping parameters (-- outFilterMultimapNmax 10, --outFilterMultimapScoreRange 1, and -- outSAMmultNmax −1). Long-read RNA-seq reads were aligned with minimap2 v2.22-r1101 using platform-specific splice-aware settings and the default secondary-alignment parameters (--secondary=yes, -N 5, -p 0.8, and -M 0.5).

For the cross-platform K562 and HepG2 analyses, raw reads from all platforms were processed as described above. ONT samples, prepared with direct RNA, direct cDNA, and cDNA_stranded library protocols, were processed individually for downstream platform-level comparisons.

For the mouse preimplantation analysis, published RNA-seq datasets of mouse oocytes and embryos from the 2-cell to blastocyst stages were used^83^.

The “2C-like system” refers to the integrated *in vivo*–*in vitro* framework used in this study to model totipotency, including 2-cell embryos, blastocysts, MERVL⁺/*Zscan4*⁺ cells (2C-like cells), MERVL⁻/*Zscan4*⁺ cells (the intermediate state between ESCs and 2C-like cells), and ESCs.

Published bulk RNA-seq datasets of human oocytes, preimplantation embryos, and hepatocellular carcinoma were analyzed using the same TEDDY workflow as described above.

BAM files generated by aligning RNA-seq reads to the corresponding reference genome were used as inputs for TEDDY. Unless otherwise specified, the mouse analyses used mm10-aligned BAM files with the GENCODE release M7 annotation and the corresponding RepeatMasker-derived mm10 TE annotation, whereas the human analyses used hg38-aligned BAM files with the GENCODE release 43 annotation and the corresponding RepeatMasker-derived hg38 TE annotation.

### TEDDY upstream analysis

Unless otherwise specified, TEDDY’s upstream processing steps in this study used the default TEDDY workflow settings. Species-matched genome builds, reference annotations, and RepeatMasker-derived TE annotations were supplied explicitly: mm10 with GENCODE M7 and mm10 TE annotation for mouse analyses, and hg38 with GENCODE 43 and hg38 TE annotation for human analyses. Input files and thread numbers were explicitly specified, whereas other reconstruction and merge parameters of the underlying StringTie engine followed default values.

### Additional transcriptomic analyses

#### Transcriptome reconstruction and harmonization

For all bulk RNA-seq datasets analyzed by TEDDY, transcriptomes were reconstructed separately for each sample from different developmental stages, cell states, or sequencing platforms as appropriate. As described above, the resulting assemblies were then processed through TEDDY’s upstream workflow, including cross-sample harmonization and annotation, TE-chimeric feature annotation, and transcript- and bin-level structural characterization and quantification. These results were organized into standard TEDDY output objects and used for downstream analyses of TE-dependent isoforms across the corresponding biological contexts.

*Transcriptome clustering.* For both the preimplantation and 2C-like systems, sample-to-sample expression correlations were first assessed by hierarchical clustering to exclude outlier replicates. Expression matrices of TE-dependent transcripts were first decomposed by non-negative matrix factorization (NMF) and then clustered by k-means to define stage- or state-associated co-expression modules.

#### Cross-species orthologous gene-level overlap analysis

Since transcript structures are not directly comparable between mouse and human, the NMF-derived stage-associated expression modules of human TE-chimeric and TE-initiated transcripts were collapsed into gene-level sets. These human gene sets were compared with mouse stage-associated TE-chimeric and TE-initiated gene sets at the mouse-human orthologous gene level, allowing cross-species recurrence to be assessed across all group-stage combinations rather than enforcing matched developmental stages.

Specifically, for each mouse developmental stage m and human expression group h, let M_m_ denote the mouse TE-associated gene set and H_h_ denote the corresponding human gene set. The overlap count was defined as *O_m_*_,ℎ_ = |*M_m_* ∩ *H*_ℎ_|, and the overlap ratio was calculated as

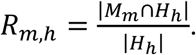

Pearson chi-square tests were performed separately for each mouse developmental stage to assess whether overlaps were evenly distributed across human expression groups. For each human group h, the test compared the number of overlapping genes with the number of mouse TE-associated genes not overlapping that group. The Pearson residual for the overlap cell was calculated as:

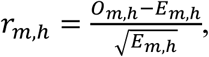

where *O_m_*_,ℎ_ and *E_m_*_,ℎ_ are the observed and expected overlap counts, respectively. Human groups with positive residuals were considered preferentially overlapping with the corresponding mouse stage, and one-sided nominal *P* values were calculated as:

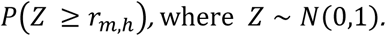

For the aggregated developmental-wave analysis, mouse stages were grouped around the 2-cell ZGA transition: oocyte was defined as the maternal wave, 2-cell as the ZGA wave, 4-cell, 8-cell, and morula as the post-ZGA wave, and blastocyst as the blastocyst wave. Human expression groups were collapsed by dominant expression pattern into maternal (Oocyte specific, Pronucleus specific, and Pre-8-cell shared), ZGA (4-cell specific and 8-cell specific), and post-ZGA (8cell/morula shared) waves. Overlap enrichment was evaluated using a one-sided hypergeometric test against a background of approximately 15,000 mouse-human orthologous genes. For each mouse wave m and human wave ℎ, with *N* background genes, K = |M_m_|, n = |H_h_|, and observed overlap *x* = |*M_m_* ∩ *H*_ℎ_|, the enrichment *P* value was calculated as the upper-tail probability under the hypergeometric distribution:

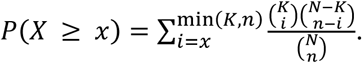

*P* values were adjusted using the Benjamini–Hochberg method across developmental-wave comparisons.

#### Single-cell RNA-seq auxiliary analysis

Single-cell RNA-seq data from the 2C-like system were processed separately for cell-state analysis^52^. These data were provided as cell-level FASTQ files and were therefore processed with the same preprocessing pipeline used for bulk RNA-seq, including STAR alignment, duplicate removal, coordinate sorting and indexing with Picard v2.26.9, and bigWig generation. Gene counts were then quantified using featureCounts v2.0.3 with GENCODE M7 (mm10). TE counts were obtained with TEcount (TEtranscripts) using the mm10 RepeatMasker-derived TE annotation. Cells with > 4,000 detected genes were retained for downstream analysis. Trajectory inference was performed using Monocle2 (DDRTree) with the top 1,500 highly variable genes selected by Seurat as ordering genes. Genes showing variation along pseudotime were identified using differentialGeneTest with a spline fit (BH FDR < 0.01).

#### HCC TE-chimeric isoform contribution and survival analysis

To evaluate the contribution of TE-chimeric isoform usage to host-gene expression, we defined an isoform-gene concordance score. For each comparison, *T*_0_ and *T*_1_ denote the mean expression of a TE-chimeric isoform in the control and case conditions, respectively. *R*_0_ and *R*_1_ denote its relative usage among all reconstructed transcripts of the host gene; and *G*_0_ and *G*_1_ denote total host-gene expression.

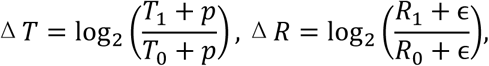

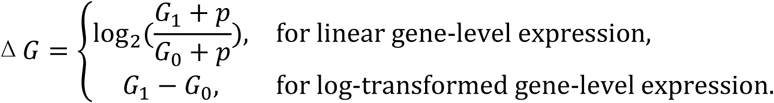

The isoform-gene concordance score was then defined as:

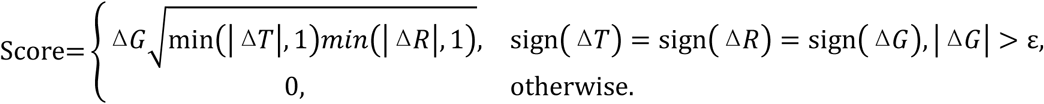

The score weights the host-gene expression change by the geometric mean of the capped isoform-expression and relative-usage changes, and is set to zero unless all three changes have the same direction. Positive and negative scores indicate concordant upregulation and downregulation, respectively. We used p = 0.3 as the expression pseudocount and ɛ = 10^−6^ to avoid division by zero. Significant TE-chimeric usage support was defined using Teddy’s chimeric-driven test with an adjusted *p* value cutoff of 0.01. Gene-level concordant support was defined as an absolute isoform-gene concordance score of at least 0.25. The score was calculated for TE-chimeric isoforms detected in three independent HCC cohorts. A host gene was considered a robust gene-level concordant candidate when its TE-chimeric isoform showed gene-level concordant support in both independent cohorts.

Enrichment of robust gene-level concordant candidates among TCGA-LIHC survival-associated genes was evaluated using Fisher’s exact test, with the background defined as genes detected in the HCC TE-chimeric analysis and also available in the TCGA-LIHC survival analysis. For survival analysis, TCGA-LIHC primary tumor samples were used, and duplicated patient-level samples were removed. Gene expression was analyzed as log_2_(FPKM + 1). Overall survival was defined using days to death when available or days to last follow-up otherwise, and vital status was used as the event indicator. Survival associations were assessed using continuous Cox proportional hazards regression.

### ChIP-seq data processing

Published Dux and Prdm14 ChIP-seq datasets were processed using an auxiliary WDL workflow. Reads were trimmed with Trimmomatic, aligned to the mm10 genome using Bowtie2, deduplicated, coordinate-sorted and indexed with Picard, and converted to RPKM-normalized bigWig tracks using bamCoverage. Peaks were called with MACS3 using matched input controls where available, with a q-value cutoff of 0.01.

### Cell culture

#### MEF and mESC culture

Mouse embryonic fibroblasts (MEFs) were derived from ICR mouse embryos at E13.5 and maintained in DMEM (Life Technologies, Cat. No. C11960500BT) supplemented with 10% (v/v) fetal bovine serum (FBS, Gibco, Cat. No. A5256701) and 1 mM L-glutamine (Merck Millipore, Cat. No. 25030164). ESCs (R1 line) were cultured on mitomycin C-treated MEFs in mESC medium containing DMEM (Merck Millipore, Cat. No. C11960500BT), 15% (v/v) FBS (HyClone, Cat. No. 16000044), 1 mM L-glutamine (Thermo Fisher Scientific, Cat. No. 25030164), 0.1 mM β-mercaptoethanol (Thermo Fisher Scientific, Cat. No. 21-985-023), 1% nonessential amino acids, and 1000 U/mL LIF (Merck Millipore, Cat. No. ESG1107). Cells were incubated at 37°C with 5% CO₂.

### Generation of ESC Lines carrying MERVL/*Zscan4* double reporters

The ESC line used for this study was R1 (mouse ES cell). The *Zscan4*-EGFP vector was generated by cloning a 2570 bp sequence upstream of the *Zscan4c* start codon^84^ into the Fugw vector (Addgene #14883) with the CMV promoter removed. The MERVL::tdTomato reporter^38^ was a gift from Samuel Pfaff (Addgene #40281). The *Zscan4*-EGFP vector reporter and the MERVL::tdTomato reporter vector were linearized and purified by QIAquick PCR Purification Kit (28106, QIAGEN). ESCs were nucleofected with 500 ng of linearized vectors in 20 μL P3 Primary Cell Nucleofector Solution (4D Nucleofector, Lonza). Stable colonies were generated by drug selection and subcloning. ESC lines containing MERVL-tdTomato and p*Zscan4c*-EGFP fluorescent reporters were validated as previously described^85^.

### Gene overexpression and knockdown in ESCs

*Nelfa* and *Arid3a* cDNAs with N-terminal HA tags were cloned into the FUW-Tet-On vector. Lentiviral particles were generated by co-transfecting the FUW-Tet-On construct, psPAX2, and pMD2.G into HEK293T cells using VigoFect. Viral supernatants were collected 48 h post-transfection and concentrated with 10% PEG 8000 at 4°C for 8–12 h. After centrifugation, the virus pellet was resuspended in 200 μL mESC medium. ESCs (∼10,000 cells/well) were infected in 96-well U-bottom plates for 8-12 h and transferred to feeder-supported ESC medium. Doxycycline hyclate (1 μg/mL) was used to induce gene expression. Overexpression efficiency was validated by qPCR and Western blotting.

### Reverse-transcription PCR and quantitative PCR

Total RNA was extracted using TRIzol reagent and reverse transcribed using 5× All-In-One RT Master Mix according to the manufacturer’s recommendations. Quantitative reverse-transcription PCR was performed with SYBR Premix Ex TaqII and the ABI7500 Fast Real-time PCR system (Applied Biosystems, Foster City, CA). The reactions were performed in triplicate using 1/10 concentration of the cDNA obtained. Relative mRNA expression was normalized to *Hprt* as an endogenous control using the ΔΔCT method.

### Embryo manipulation, microinjection, and *in vitro* culture

#### Embryo collection

Specific pathogen–free C57BL/6J female mice (8–10 weeks) were superovulated by injection of 7 IU PMSG, followed 48 h later by 5 IU hCG. MII oocytes were collected from the oviducts of unmated females. For zygote collection, PN3-stage embryos were isolated based on pronuclear morphology. All animal procedures were approved by the Biological Research Ethics Committee of Tongji University.

#### siRNA microinjection

For maternal knockdown, siRNAs targeting *Arid3a* were diluted to a working concentration of 5 μM in nuclease-free water. Approximately 10 pL of siRNA solution was injected into the cytoplasm of MII oocytes using a Piezo-driven micromanipulator. Injected oocytes were allowed to recover for ≥1 h prior to fertilization.

#### Intracytoplasmic sperm injection (ICSI)

ICSI was performed as previously described, using HEPES-buffered CZB (HCZB) for gamete handling. Sperm heads were injected into siRNA-injected oocytes, and successfully fertilized embryos were transferred to CZB medium under mineral oil and cultured in 5% CO₂.

#### Embryo culture and developmental assessment

Embryos were monitored from the 2-cell stage to the blastocyst stage. Developmental progression was quantified by scoring embryos at each cleavage stage. Control embryos progressed to blastocysts at nearly 100%, whereas *Arid3a*-depleted embryos predominantly arrested at the morula stage.

#### Long-read RNA sequencing of 2-cell embryos

Total RNA quality and quantity were assessed using a NanoDrop One spectrophotometer, a Qubit 4.0 Fluorometer, and an Agilent 2100 Bioanalyzer. Samples with an RNA integrity number greater than 8.0 were used for library preparation. Poly(A)+ RNA was isolated using the Dynabeads™ mRNA Purification Kit, and cDNA-PCR libraries were prepared using the Oxford Nanopore cDNA-PCR Sequencing Kit V14 (SQK-PCS114) according to the manufacturer’s protocol. Libraries were sequenced on a PromethION platform using FLO-PRO114M flow cells at GrandOmics (Wuhan, China).

## Acknowledgements

We thank our colleagues in the lab for their help with experiments and manuscripts. We specifically thank Wenqiang Liu and Yifan Sheng for their assistance with embryo injections, and Jiatong Sun for help with RNA library construction.

## Author contributions

Y.X., R.L. and S.G. conceived and designed the study. Y.X. developed the computational framework. L.S. performed most of the experiments. Y.L., J.Y. and H.W. assisted with the experiments. Y.X. designed and performed the data analyses. Y.X. and R.L. interpreted the findings. Y.X. wrote the manuscript. J. S and R.L. contributed to manuscript revision. C.J. and Y.Z. reviewed the manuscript. S.G. supervised the project.

## Data availability

All sequencing data supporting this study are available from public repositories or have been newly deposited. Newly generated sequencing data have been deposited in the Genome Sequence Archive (GSA), including H3K4me3 ChIP-seq and RNA-seq data from mouse embryonic stem cells and 2C-like cells (PRJCA068954) and long-read RNA-seq data from mouse 2-cell embryos (CRA046285). Publicly available data used in this study include mouse preimplantation RNA-seq (GSE97778)^83^, Dux ChIP-seq (GSE95517)^51^, Prdm14 ChIP-seq (GSE25409)^86^, H3K27ac and H3K4me3 ChIP-seq from mouse preimplantation embryos and ESC subpopulations (GSE71434, GSE73952 and GSE164486)^85,87,88^, hepatocellular carcinoma RNA-seq (GSE101432, GSE135631 and GSE214846)^67,68^, human early embryonic development RNA-seq (GSE44183)^58^, and single-cell RNA-seq (E-MTAB-5058) from ArrayExpress; TCGA HCC data accessed via the NCI Genomic Data Commons. For benchmark comparisons, we used the *Drosophila* dataset from the ChimeraTE repository (https://github.com/OliveiraDS-hub/ChimeraTE) and matched multi-platform sequencing data (Illumina, PacBio, Oxford Nanopore) for K562 and HepG2 cell lines from the SG-NEx project^32^.

## Code availability

The custom code for the TEDDY pipeline is available on GitHub: https://github.com/YeehanXiao/Teddy. Benchmarking scripts and the configuration files are available at https://github.com/YeehanXiao/TEDDY-reproducibility.

