## Supplementary Information for "TEDDY: An integrative workflow for TE-chimeric isoform reconstruction and systematic characterization of TE-dependent transcriptional regulation"

### Supplementary Figures:

#### Fig. S1 | Locus-level resolution and validation across simulated datasets

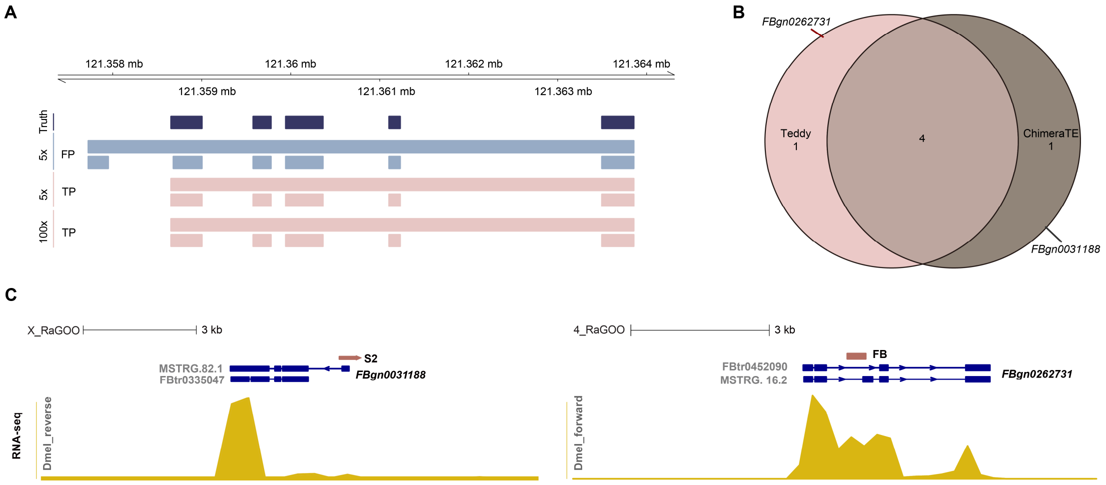


**A**, Resolution of isoform structure with increasing sequencing depth. At low coverage (5×), TEDDY assembles two isoforms: one true positive (TP) matching the ground truth and one false positive (FP) containing an extra exon. At higher coverage levels (10×, 25×, 50×, and 100×), the TP isoform is consistently recovered while the FP isoform is excluded, with 100× shown as a representative high-coverage example.

**B**, Venn diagrams summarizing overlap of TE-chimeric calls between TEDDY and ChimeraTE. The tools shared four calls in the *Drosophila* dataset; each made one unique prediction that was manually inspected and is shown in Fig. S1C.

**C**, Manual inspection of unique calls. (Left) A false positive call from ChimeraTE at the *FBgn0031188* locus, driven by a mis-oriented S2 element. (Right) A true positive FB-chimeric isoform reported by TEDDY at the *FBgn0262731* locus, with strand-concordant coverage support.

**Fig. S2 | Examples of TE-Chimeric transcript structures in mouse oocytes**

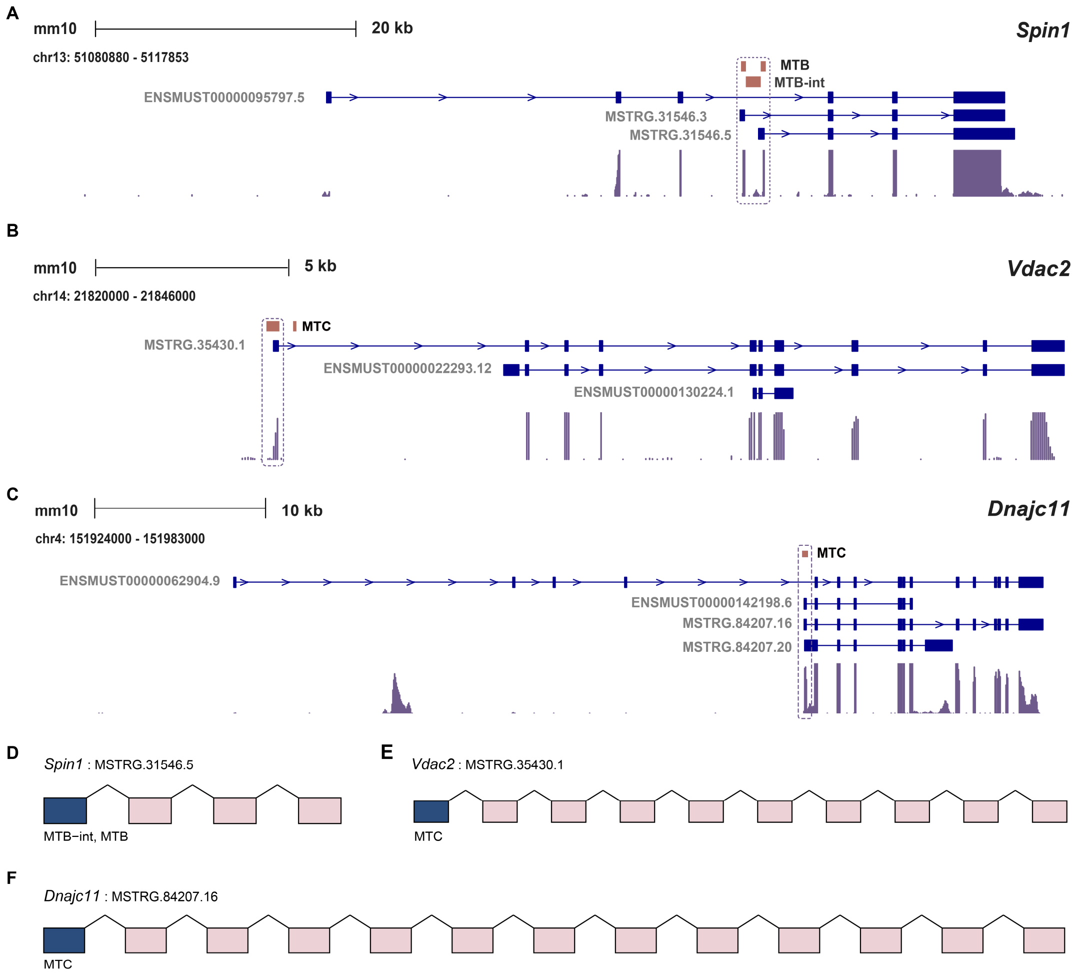


**A**, Structures of *Spin1* isoforms in mouse oocytes. Based on the GENCODE (Release M7) annotated mouse genome, TEDDY assembled both the canonical (ENSMUST00000095797.5) and the truncated TE-chimeric isoforms of *Spin1* (MSTRG.31546.3 and MSTRG.31546.5) initiated from MTB or MTB-int. The RNA-seq track shows the relative abundance of RNA at the exon level, visualized using BigWig files from processed RNA-seq data. Names of the novel transcripts were assigned by TEDDY during transcript assembly and may vary with each de novo GTF generation. Therefore, the names of TE-chimeric transcripts in Fig. S2 only serve for reference and distinction purposes, and this applies to the subsequent analysis hereafter. **B**, Transcript structure of MTB-initiated *Spin1* TE-chimeric isoform generated by TEDDY. Blue boxes: TE-derived alternative exons. Pink boxes: canonical exons.

**C**, Structures of *Vdac2* isoforms in mouse oocytes. TEDDY detected an elongated MTC-initiated chimeric isoform (MSTRG.35430.1) and the canonical transcript (ENSMUST00000229232.12).

**D**, Transcript structure of the MTC-initiated elongated chimeric isoform (MSTRG.35430.1) for the gene *Vdac2* identified by TEDDY. Blue boxes: TE-derived alternative exons. Pink boxes: canonical exons.

**E**, Structures of *Dnajc11* isoforms in mouse oocytes. TEDDY has identified novel TE-chimeric isoforms such as MSTRG.84207.16 and MSTRG.84207.20 alongside the canonical isoforms, such as ENSMUST00000142198.8, which is documented in GENCODE.

**F**, Transcript structure of a newly assembled MTC-initiated isoform for *Dnajc11* (MSTRG.84207.16) identified by TEDDY.

**Fig. S3 | Gene expression analysis in mouse preimplantation embryos and ESCs**

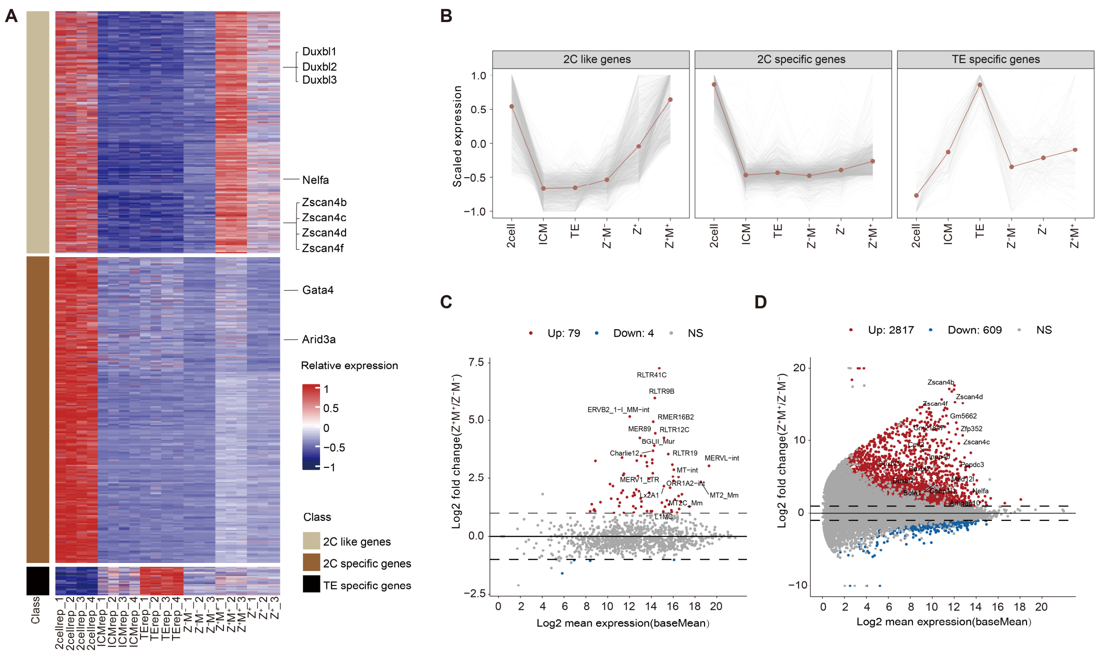


**A**, Heatmap generated from cluster analysis of gene expression in ESCs, MERVL$^+$/*Zscan4*$^+$ cells, *Zscan4*$^+$ cells, 2C embryos and blastocysts (separated inner cell mass (ICM) and trophectoderm (TE)) using degPattern analysis by DEGreport. Three distinct groups of genes were categorized based on their expression patterns. Each row represents the Z score of expression level (FPKM). Z$^-$M$^-$: ESCs; Z$^+$: *Zscan4*$^+$ cells (intermediate cell state in the transition between ESCs and 2C-like cells); Z$^+$M$^+$: 2C-like cells.

**B**, Line charts depicting the gene expression patterns in mouse preimplantation embryos for the three gene clusters identified in S3A. Z$^-$M$^-$: ESCs; Z$^+$: *Zscan4*$^+$ cells (intermediate cell state in the transition between ESCs and 2C-like cells); Z$^+$M$^+$: 2C-like cells.

**C**, Volcano plot showing differentially expressed TEs between 2C-like cells (MERVL$^+$/*Zscan4*$^+$ cells) and ESCs. Red dots represent upregulated TEs, and blue dots represent downregulated TEs in 2C-like cells compared with ESCs (TE subfamilies with FPKM change greater than 2-fold or less than 0.5-fold). n = 3 biological replicates.

**D**, Volcano plot displays differentially expressed genes between 2C-like cells (MERVL$^{+}$/*Zscan4*$^{+}$ cells) and ESCs. Red dots represent upregulated genes, and blue dots represent downregulated genes in 2C-like cells compared with ESCs (Genes with FPKM change greater than 2-fold or less than 0.5-fold). n = 3 biological replicates.

**Fig. S4 | MT2-chimeric transcripts are enriched in 2C-like cells**

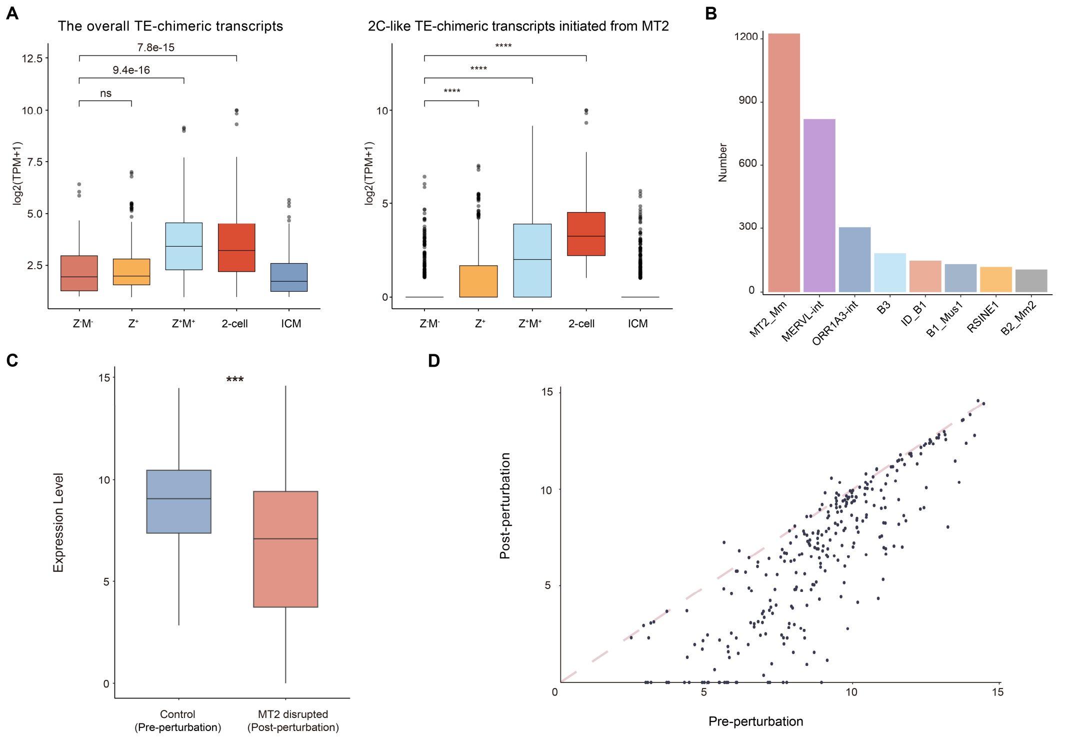


**A**, Boxplot showing the expression levels of TE-chimeric transcripts in the indicated cells and 2-cell embryos. Left: expression levels of all TE-chimeric transcripts identified in the indicated cells, ICM and 2-cell embryos. Right: expression levels of 2C-like TE-chimeric transcripts associated with MT2 in the indicated cells, ICM and 2-cell embryos. The Wilcoxon rank-sum test (Mann-Whitney U test) was used. $Z^-M^-$: ESCs; $Z^+$: *Zscan4*$^+$ cells (intermediate cell state in the transition between ESCs and 2C-like cells); $Z^+M^+$: 2C-like cells.

**B**, The bar graph showing the number of top transposable elements (TEs) associated with shared TE-chimeric transcripts between 2-cell embryos and 2C-like cells.

**C**, Genes carrying significant MT2_Mm-driven chimeric isoforms identified by TEDDY are downregulated following CRISPRi-mediated repression of MT2 (MT2i) in mouse 2-cell embryos compared to control (CTRi), using published RNA-seq data[1]. The center of the box plots represents the median value, and the lower and upper lines represent the 25% and 75% quantiles, respectively. Statistical significance was assessed using a two-sided *t*-test on log-transformed counts.

**D**, This scatter plot shows the expression change of genes with significant MT2_Mm-driven chimeric isoforms identified by TEDDY in MT2i 2-cell embryos compared with control 2-cell embryos.

### Fig. S5 | Experimental validation of TEDDY-predicted TE-chimeric isoforms

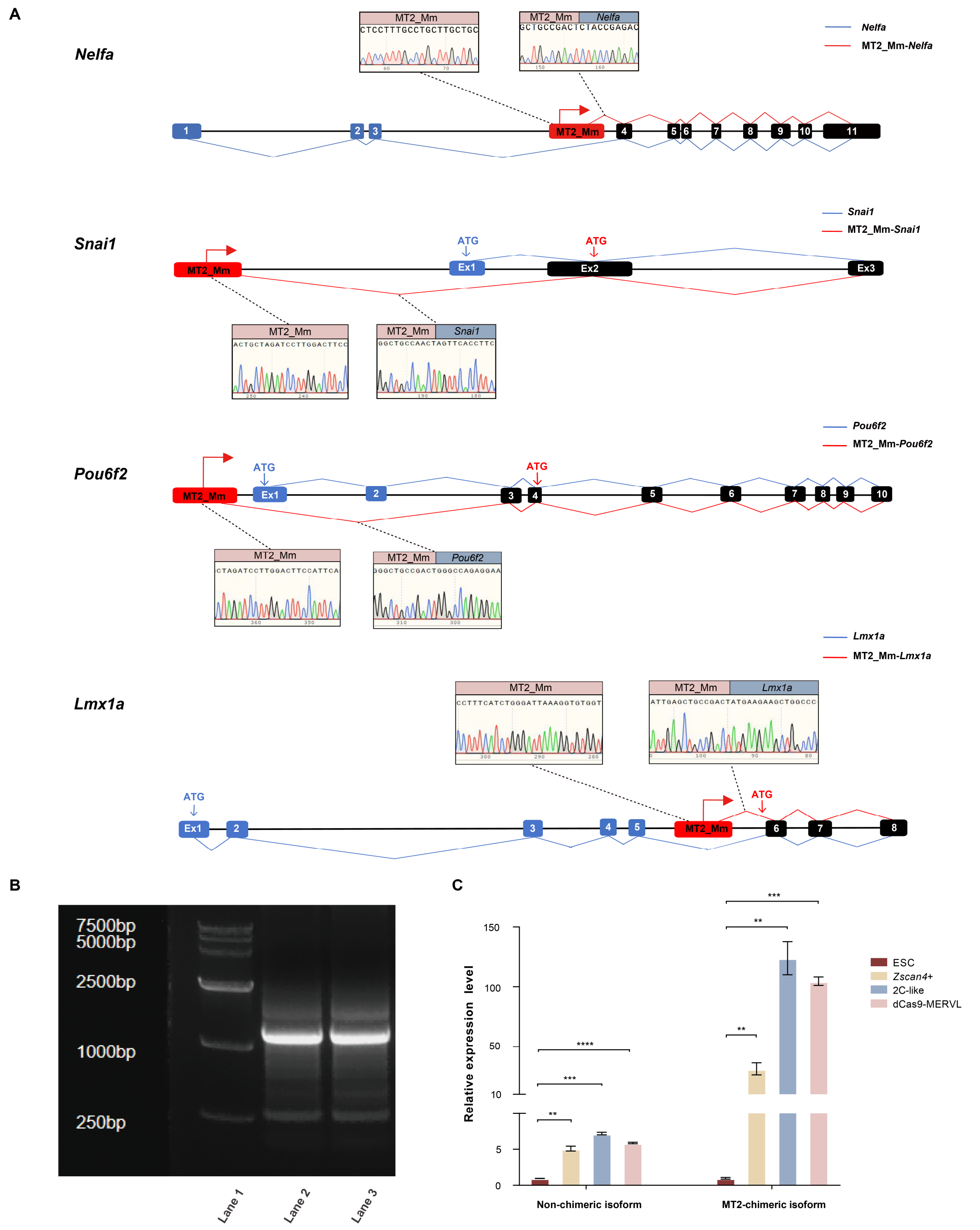


**A**, Schematic representations of the canonical (non-chimeric) and TEDDY-predicted TE-chimeric isoforms for *Nelfa*, *Snai1*, *Pou6f2*, and *Lmx1a*, arranged top to bottom. For each locus, adjacent Sanger sequencing chromatograms of the RT-PCR products confirm the TE–host splice junctions captured by TEDDY.

**B**, RT-PCR validation of the chimeric isoform of *Nelfa* in 2C-like cells. Lane 1, DNA ladder; lanes 2–3, RT-PCR products from 2C-like cells. The expected amplicon size is approximately 1437 bp.

**C**, qRT-PCR showing expression levels of the canonical non-chimeric isoform and the TE-chimeric isoform of *Nelfa* in ESCs, *Zscan4*$^+$ cells, 2C-like cells and CRISPRa-induced 2C-like cells (2C-like cells induced by CRISPR-mediated activation of MERVL)[2]. Shown is mean ± SD; n = 3 biological replicates. Significance was analyzed using Student's *t*-test. $*p < 0.05$, $**p < 0.01$, $***p < 0.001$, $****p < 0.0001$.

**Fig. S6 | Transcription Factors associated with 2C TE-initiated chimeric transcripts predicted by TEDDY**

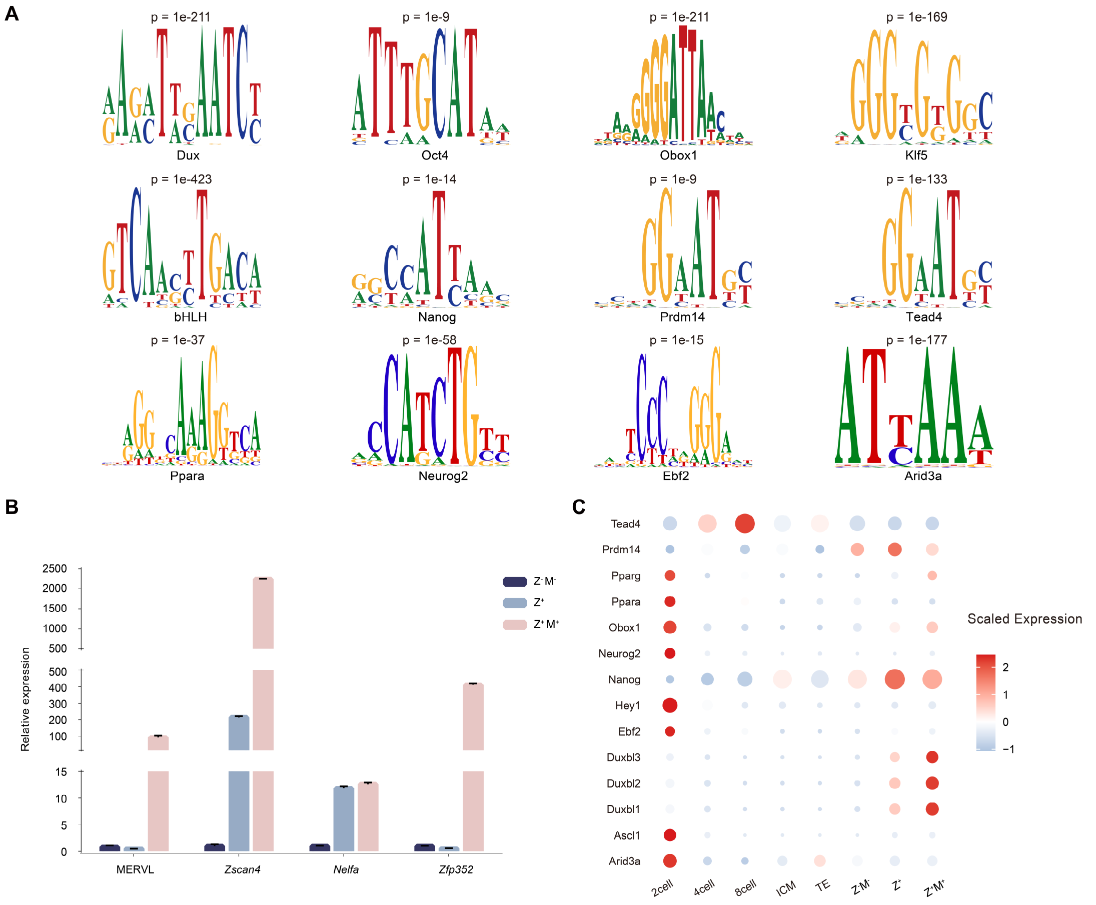


**A**, TF binding motifs enriched at TEs linked to 2C-like TE-chimeric transcripts. Representative motifs are shown; Homer-reported p-values are indicated above each motif.

**B**, Bar plot showing the expression levels of crucial 2C markers, including *Zscan4* and MERVL, in ESCs ($Z^-M^-$), *Zscan4*$^+$ cells ($Z^+$), and 2C-like cells ($Z^+M^+$).

**C**, Dot plot depicting scaled expression levels of transcription factors in the indicated developmental stages of the mouse early embryos and the indicated cells, as determined by RNA-seq analysis. The size of the dot reflects the relative expression magnitude.

#### Fig. S7 | Characterization of Dux Binding Sites in ESCs

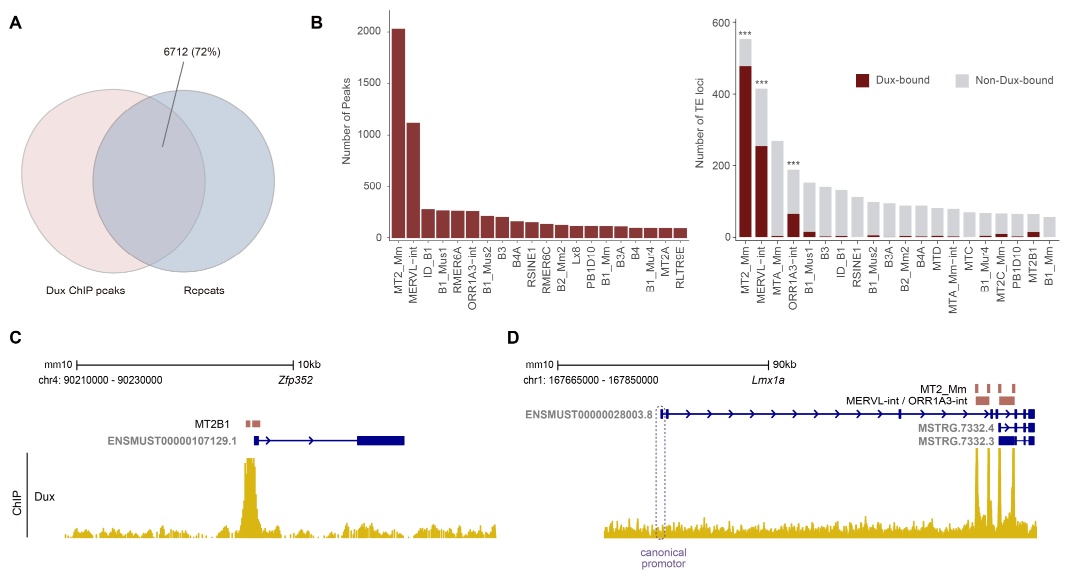


**A**, Venn diagram depicting the overlap between Dux ChIP-seq peaks in ESCs and repetitive sequences, revealing that 72% of Dux binding sites (6712 peaks) are located within repetitive elements.

**B**, Distribution of Dux peaks across repetitive elements. The left panel summarizes the subfamily composition of all Dux-bound repeats. The right panel shows TE loci associated with 2C-specific and 2C-like TE-chimeric transcripts, stratified by Dux peak overlap. Enrichment of Dux binding for each TE subfamily, relative to all other TE-chimeric-associated loci, was evaluated using a one-sided Fisher's exact test with Benjamini-Hochberg correction (***, adjusted $P < 0.001$), revealing a significant enrichment of MT2-associated elements.

**C**, Transcript structures of *Zfp352* and the genomic track of Dux binding signal at the *Zfp352* locus. This particular isoform is currently included in the GENCODE (Release M7), supporting the reliability of TEDDY in detecting TE-chimeric transcripts.

**D**, Transcript structures of *Lmx1a* and the genomic track of Dux binding signal at the *Lmx1a* gene locus. Both the canonical and newly identified truncated TE-initiated isoforms of *Lmx1a* were captured by TEDDY.

**Fig. S8 | Single-Cell Pseudotime Analysis of 2C-like Transition**

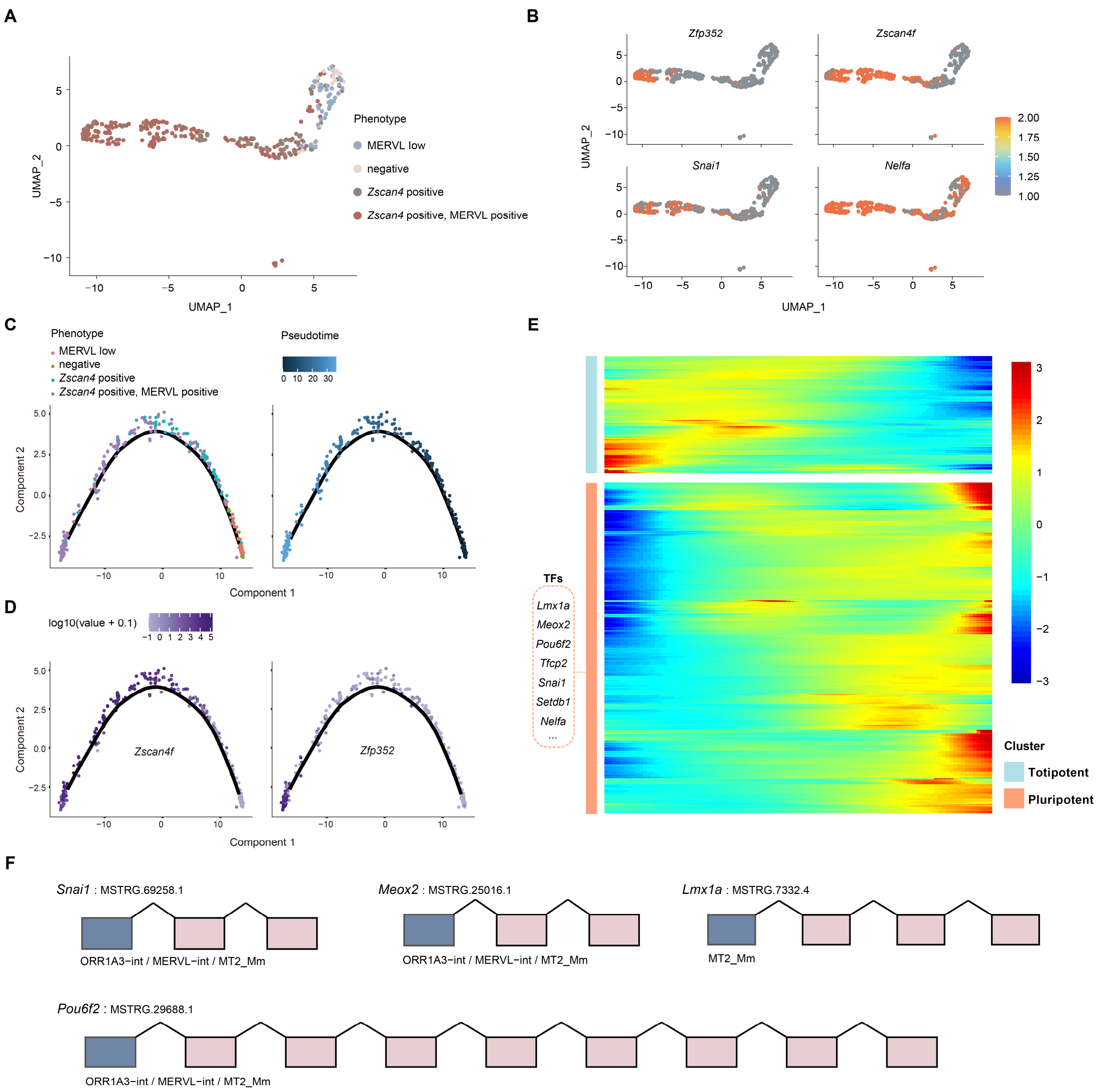


**A**, UMAP analysis revealing a continuum of transcription states of cells undergoing 2C-like conversion. Dots on the UMAP plot represent individual cells, colored according to their FACS-derived phenotypic states from the original study (based on the expression of 2C reporters *Zscan4* and MERVL).

**B**, UMAP plots showing the expression patterns of the 2C markers along the pseudotemporal trajectory.

**C**, Pseudotemporal trajectory analysis showing the relative progression of each cell in the 2C-like state transition. Each dot represents a single cell.

**D**, Expression patterns for the 2C markers *Zscan4f* and *Zfp352* along the pseudotemporal trajectory of 2C-like transitions. Colors represent the Z score of log2-transformed FPKM values.

**E**, Heatmap showing expression changes of TFs along a pseudo-temporal trajectory (horizontally) in the transition between ESCs and 2C-like cells. The color scale indicates scaled relative expression levels, with warmer colors representing higher relative expression and cooler colors representing lower relative expression. Totipotent-associated TFs (blue) and pluripotent-associated TFs (red) are defined based on their expression levels in ESCs and 2C-like cells. Representative totipotent-associated TFs bearing TE-initiated transcripts are labeled on the left, with the structure information detailed in Table 3.

**F**, Transcript structures of TE-chimeric isoforms for totipotency-related TFs including *Snai1*, *Meox2*, *Lmx1a*, and *Pou6f2*.

**Fig. S9 | *In Vitro* and *In Vivo* Validation of ARID3A in 2C-like Transition**

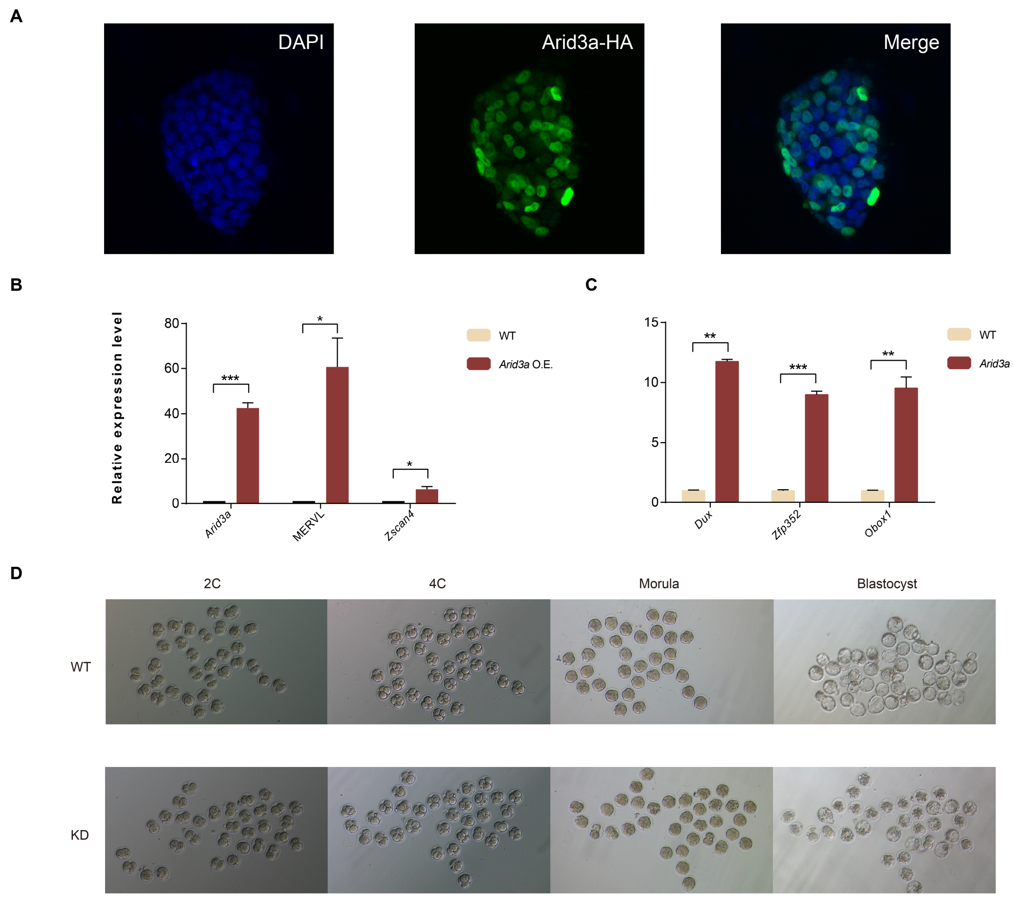


**A**, Immunofluorescence staining showing the nuclear localization of Arid3a in *Arid3a* O.E. ESCs. Arid3a expression was visualized with GFP. Cell nuclei were visualized with DAPI. Representative of three independent experiments.

**B-C**, RT-qPCR analysis showing elevated expression of 2C-associated markers (MERVL, *Zscan4*, *Dux*, *Zfp352*, *Obox1*) in *Arid3a*-overexpressing ESCs compared to WT controls. Gene expression was normalized to *Hprt*. Statistical significance was assessed by unpaired *t*-test with Welch's correction, *$p < 0.05$, **$p < 0.01$, ***$p < 0.001$. Three independent experiments.

**D**, Representative images showing the different stages of WT and *Arid3a* KD mouse embryos. Representative of two independent experiments.

**Supplementary Table 1 | Benchmarking and Comparison of Representative TE-related Transcript Analysis Tools**

This table summarizes tool scope, benchmark scoring strategy, raw accuracy metrics, read-level support for simulated TE-host breakpoint structures, and measurements of runtime and memory usage.

Sheet 1 summarizes the primary goals, methodological strategies, platform support, downstream analysis support, and software implementation characteristics of related tools discussed in this study (Supplementary Note 1). Tools with an asterisk are listed for contextual comparison as TE quantification methods and were therefore excluded from the TE-chimeric benchmark.

Sheet 2 describes the tool-matched scoring strategy used to harmonize benchmark outputs at the host-gene level. All tools were evaluated using the same simulated datasets, while the ground-truth set and positive criterion were matched to each tool's native settings and intended output.

Sheet 3 reports the raw benchmark accuracy metrics underlying Fig. 2A across sequencing depths. Predicted Genes and Truth Genes indicate the numbers of predicted host genes and tool-compatible truth genes defined according to the scoring strategy in Sheet 2. TP, FP, and FN were then derived from these gene sets, and precision, recall, and F1 were calculated from TP, FP, and FN.

Sheet 4 reports read-level support for simulated TE-chimeric breakpoint structures. Loci were grouped according to the length and local topology of the TE-incorporated segment. Short TE full-span events were defined as TE-overlap segments shorter than 125 bp and embedded within an exon, such that the TE-derived segment and adjacent host sequence could be spanned by individual sequencing reads. Long TE boundary events were defined as TE-overlap segments of at least 125 bp. The upper section summarizes expressed loci across sequencing depths, whereas the lower section summarizes 100x loci stratified by TPM group. TPM groups were defined using transcript abundance values pre-assigned in the simulation. Read-level support was assessed using anchor-based and continuous-alignment evidence; N. AnchorPair Supported and N. Continuous-read Supported report loci supported by each strategy, whereas N. Supported reports their union (see Methods). Support Rate was calculated as N. Supported divided by N. Truth Loci.

Sheets 5 and 6 report runtime and memory usage on the 100x simulated dataset and the matched *Drosophila* demonstration dataset, respectively. Wall-clock time and peak resident memory were measured across three independent runs under the same computational environment.

**Supplementary Table 2 | Downsampling-based Recovery of TE-chimeric Transcripts in Real K562 and HepG2 Datasets**

Real K562 and HepG2 Illumina and ONT cDNA datasets were downsampled to 10%, 25%, 50%, and 75% of the original reads, and each downsampled dataset was processed through the TEDDY workflow.

Sheet 1 summarizes within-platform recovery of full-depth TE-chimeric isoforms using the corresponding full-depth transcriptome from the same platform as reference.

Sheet 2 summarizes recovery of TE-chimeric isoforms jointly supported by Illumina and ONT in the cross-platform analysis. Exact recovery denotes concordance of transcript structure between downsampled and full-depth transcriptomes.

Sheet 3 summarizes structure-coupled quantification concordance across downsampling depths. For exact-matched TE-chimeric transcripts with joint structural support from both platforms, we retained those with $\text{FPKM} \geq 1$ in the original full-depth Illumina dataset. Pearson and Spearman correlations were then calculated between $\log_2$(FPKM+1) values from independently reconstructed and quantified downsampled Illumina datasets and the corresponding full-depth estimates.

**Supplementary Table 3 | TE Subfamily Composition of TE-chimeric Transcripts in Mouse Preimplantation Developmental Expression Groups**

This table summarizes the proportions of major TE subfamilies associated with TE-chimeric transcripts in the developmental expression groups identified in Fig. 3A. Sheet 1 provides a summary of the nine expression groups, including group names, expression features, the number of TE-chimeric transcripts, and major TE categories. Subsequent sheets report the TE category and subfamily composition for each group or related group set. Minor and unclassified TE subfamilies were grouped as "Others" for readability.

**Supplementary Table 4 | Annotated 2C-like TE-initiated Transcripts Identified by TEDDY**

This table lists 2C-like TE-initiated transcripts identified by TEDDY using the GENCODE M7 annotation, including genomic coordinates, transcript IDs, host genes, associated TE names, and TE classes. Cross-version stability is indicated by Teddy_M25_reconstructed (whether the transcript remains reconstructable by TEDDY using GENCODE M25) and Catalog_M25 (inclusion in the official GENCODE M25 catalog). Nanopore read-level support from 2-cell embryo data is provided by Nanopore_JunctionSupported (TE-host junction support) and Nanopore_ExonchainSupported (exon-chain structural support).

**Supplementary Table 5 | Primers for RT-PCR amplification and Sanger validation of TEDDY-predicted TE-chimeric isoforms**

This table lists the forward and reverse primers used to amplify TEDDY-predicted TE-chimeric isoforms validated in Fig. S5. PCR products were further confirmed by Sanger sequencing.

**Supplementary Table 6 | TE Category Composition and Abundance Comparison in Human Preimplantation Embryos**

This table summarizes the TE categories shown in Fig. 6B and their proportions across human preimplantation developmental groups. Sheet 1 lists each TE category and the TE subfamilies included in that category. The remaining columns show the percentage (%) of TE-chimeric transcripts associated with each TE category in each developmental group. Minor and unclassified TE subfamilies were grouped as "Others" for readability. Sheet 2 summarizes the relationship between 8-cell-specific TE-chimeric transcripts and subfamily-level TE abundance comparisons between 8-cell embryos and adjacent developmental stages, with the Rate (%) indicating the proportion of TE-chimeric events in each category. For the TE subfamilies highlighted in the differential abundance analysis (Fig. 6C), Sheet 3 details whether specific TE copies are located within gene bodies, whether the corresponding host genes are expressed at the 8-cell stage, and whether associated TE-chimeric transcripts are detected at these loci. No. Gene-body TE Copies indicates TEs within annotated genes (Host Genes), and No. Expressed Host Genes in 8C counts the subset expressed at the 8-cell stage (mean $\mathrm{CPM} \geq 1$). Associated TE-chimeric genes indicate the number of gene loci producing TE-chimeric transcripts involving the indicated TE subfamily; TE-chimeric genes expressed in 8C indicate the subset of these loci with associated TE-chimeric transcripts expressed at the 8-cell stage (mean $\mathrm{FPKM} > 1$).

**Supplementary Table 7 | Ortholog-level assessment of mouse oocyte-associated TE-chimeric candidates in human preimplantation embryos**

This table lists representative mouse-human orthologous gene pairs with TE-initiated isoforms detected in mouse oocyte-associated groups and human preimplantation embryos, examined as examples of conserved or recurrent TE-associated activation. Mouse isoforms were selected from the expression clusters Oocyte, Oocyte_2cell, and Shared group1, representing TE-initiated transcripts expressed in oocytes. Human isoforms were required to have $\mathrm{FPKM} \geq 5$ in at least one human preimplantation stage. The table lists gene symbols, Ensembl gene IDs, TE subfamilies, and human stage-wise $\mathrm{FPKM}$ values.

#### Supplementary Note 1. Detailed technical comparison of related tools for TE-chimeric transcript analysis

##### Biological scope and structural inference strategies

Early tools such as SQuIRE, TEtranscripts, and Telescope focused solely on locus- or subfamily-level TE expression. The growing recognition of TE-driven promoters as *cis-regulatory* elements then motivated the development of methods designed to detect TE-initiated transcripts, including LIONS[3] and TEProf2[4]. Subsequent work further extended the analytical focus to internal TE exonization (ChimeraTE)[5] and protein-coding potential (FREDY)[6]. These methods differ in focus and thus in their strategies, as detailed below. Although we also included Arriba[7] in parts of the benchmarking analysis, we did so to assess how a widely used gene-fusion detection strategy generalizes to TE-associated chimeric events. As Arriba was not designed for TE-associated transcript analysis, it was not included in Supplementary Table S1.

Related tools for TE-chimeric transcript analysis differ fundamentally in their transcript inference strategies, beginning with how candidate TE-associated transcripts are defined, reconstructed, and represented. A major point of divergence lies in the intended output of these methods, which determines whether transcriptome reconstruction is required as a prerequisite. Event-focused tools like ChimeraTE and LIONS are designed to report TE-associated events or chimeric junctions and therefore do not recover full-length transcript structures. ChimeraTE's Mode 1 infers chimeric transcripts based on the genomic proximity of separately aligned gene and TE reads, an approach that is vulnerable to mapping biases in repetitive regions and incapable of resolving complex splicing patterns[3,5]. Mode 2 uses Trinity for *de novo* contig assembly, and contigs containing TE-derived sequence are subsequently matched to reference transcripts based on sequence similarity. This mode operates at the contig-assembly level and does not provide genome-coordinate exon-chain reconstruction. Transcriptome-wide *de novo* assembly is also computationally demanding in large, repeat-rich mammalian genomes, limiting its scalability for systematic TE-chimeric isoform analysis. Similarly, although LIONS includes a Cufflinks-based *de novo* assembly step, its final output remains event-level and does not retain exon structure.

Recently, workflow-oriented approaches have emerged, reflecting a growing consensus that transcriptome reconstruction is essential to systematically identify TE-dependent isoforms [8]. Transcriptome analysis generally relies on three distinct paradigms: pure de novo assembly (e.g., Trinity), strictly annotation-dependent methods, and reference-guided reconstruction (e.g., Cufflinks, StringTie). For TE-chimeric transcripts, reference-guided reconstruction is particularly suited. While TE-chimeric isoforms originate in repeat-rich sequences, their functional interpretation demands full host-gene context. Contig-based *de novo* assemblies lack genome-coordinate exon-chain resolution, whereas strict annotation-dependent approaches miss unannotated TE-chimeric isoforms. Reference-guided reconstruction therefore provides a practical balance: it preserves precise genomic anchoring to resolve exon structures, while enabling the recovery of novel transcripts through local *de novo* discovery. Within this strategy, StringTie/StringTie2 provides a particularly suitable foundation because it models each locus as an alternative splicing graph, using the local read-coverage distribution and splice-junction support to infer transcript paths under a flow-network framework. This StringTie-based strategy is shared by TEProf2, FREDY, and TEDDY, but the implementation differs substantially. TEProf2 is distributed as a collection of analysis scripts rather than as a cohesive software package, whereas FREDY provides a modular command-line workflow around external tools. TEDDY integrates bundled StringTie executables into R package functions for parallel reconstruction across samples. Previous studies, along with our own findings, highlight the regulatory importance of context-specific TE-chimeric transcripts.

#### Cross-sample transcriptome consolidation

To compare isoform usage across samples, independently assembled transcripts must first be merged into a single, non-redundant reference. However, this process is frequently confounded by sample-specific identifiers, overlapping isoforms, and minor boundary shifts—even among transcripts with identical exon chains. Current methods fail to resolve this issue at the exon-chain level. For example, while TEProf2 annotates candidate transcripts against the reference, this remains based on genomic-range overlap rather than matching precise exon-intron topologies. To overcome this limitation, TEDDY directly aligns and merges transcripts across samples based on their structural boundaries. Through systematic filtering and annotation, TEDDY generates a unified transcriptome reference.

#### Dual-resolution quantification: transcript inference and bin-level counting

RNA-seq quantification can be performed at different feature resolutions, including gene-level, transcript-level, and local exon/bin-level measurements. Gene-level quantification is relatively straightforward because reads mapped to the same gene can usually be pooled without resolving their transcript of origin. For short-read RNA-seq, multiple isoforms of the same gene often share exons and splice junctions, so a read may be compatible with more than one transcript. Long-read data can also be affected by splice-site jitter, coverage variation, and sequencing-depth limitations. Thus, transcript-level quantification is inherently an inference problem rather than a direct counting problem. To address this mapping ambiguity, graph-based methods like StringTie iteratively allocate read flow across compatible paths using residual coverage, meaning transcript-level estimates are model-based inferences rather than simple arithmetic partitions of gene-level counts. LIONS calculates RPKM from normalized coverage over the genomic range of each transcript. This provides expression support for TE-initiated transcripts based on read coverage across the assigned interval, but does not model ambiguous read assignment among alternative isoforms. FREDY and TEProf2 instead adopt StringTie-derived abundance estimates from isolated single-sample reconstructions as their transcript-level abundance values. In contrast, TEDDY first consolidates sample-level reconstructions into a unified, non-redundant transcriptome reference and then re-quantifies each sample against this shared structure. With candidate transcript structures fixed as a topological prior, TEDDY reuses the same flow-network inference framework to assign read abundance to each isoform within the unified reference space. This re-quantification is implemented through the TEDDY::stringtieCombine() interface. By invoking StringTie in -e mode, this function disables *de novo* structural assembly and restricts abundance estimation to the unified reference, thereby ensuring the quantification is coupled to the consensus structure. The resulting coverage, FPKM, and TPM values are extracted from the re-quantified GTF files and organized into a transcript-by-sample abundance matrix stored as a SummarizedExperiment object.

Transcript-level abundance estimates describe the expression of reconstructed full-length isoforms, whereas evaluating local alternative splicing events, such as TE-driven exon inclusion, alternative promoter usage, or local exon replacement, requires an additional local resolution often lacking in existing TE-chimeric tools. Context-specific TE-chimeric events can be reflected by changes in the local read distribution across multiple samples. TEDDY therefore incorporates a bin-level quantification strategy to capture local TE-associated exon usage. Overlapping transcript structures are flattened into a comprehensive set of mutually exclusive exon-derived intervals, hereafter referred to as bins (Fig. 1A). Sequencing reads are then assigned directly to these non-overlapping regions to calculate absolute local read counts. TEDDY wraps GenomicFeatures for exon partitioning and bin-level read counting. Together, these two layers distinguish the abundance of reconstructed isoforms from the local usage of TE-associated exon bins, allowing TEDDY to analyze TE-dependent isoform remodeling.

**Software architecture and reproducibility**

The software organization of these tools also affects usability, reproducibility, and scalability. As summarized in Supplementary Table S1, several existing methods are distributed as heterogeneous script-orchestrated workflows combining R, Python, shell scripts, and external command-line tools. Such designs often require manual dependency management, intermediate-file handling, and execution across different software environments; in some cases, they also depend on older software versions or tools that are no longer actively maintained, such as Cufflinks. TEDDY instead provides an end-to-end R-based analysis environment that reduces external dependency configuration while making the roles of integrated tools explicit. It further connects transcriptome reconstruction, TE annotation, transcript- and bin-level quantification, isoform-usage analysis, visualization, and regulatory interpretation within a single R/Bioconductor-compatible workflow.

**Additional validation of simulated benchmark data**

Simulated FASTQ files were re-quantified against the full ground-truth transcriptome with RSEM to confirm overall agreement with the pre-assigned TPM values. We further examined structural support in the genome-coordinate BAM files. Breakpoint-informative support increased with sequencing depth and pre-assigned transcript abundance. At 100×, 90.30% of expressed long TE boundary loci and 94.51% of expressed short TE full-span loci were supported, and exact CIGAR-N split-read support was observed at 147 of 186 unique TE-host splice junctions from expressed loci (79.03%). These results support the quantitative and structural fidelity of the simulated reads and indicate that the lower performance of junction- or event-based tools is unlikely to reflect a general lack of appropriately aligned supporting reads (Supplementary Table 1).

**Computational performance analysis**

We evaluated runtime and memory usage for the 100× simulated benchmark dataset, using the same inputs and tool configurations as in the accuracy benchmark (Supplementary Table 1). Each tool was run across three independent runs, and wall-clock time and peak RSS were recorded at the system level using the Linux /usr/bin/time -v command. Peak RSS refers to peak resident set size. In this benchmark setting, TEDDY completed the full workflow with moderate runtime and memory usage.

Following the locus-level comparison shown in Fig. S1C, we used the same single-sample *Drosophila* demonstration dataset to evaluate runtime and memory usage for a matched TE-chimeric identification task. Both tools were run on the same input data across three independent runs on a Linux server equipped with two AMD EPYC 7763 64-core processors, 256 CPU threads, and 881 GiB total memory. Wall-clock time and peak resident memory were recorded for each run at the system level using the Linux /usr/bin/time -v command. ChimeraTE was run using its Mode 1 example command with --strand rf-stranded.

In this single-sample, single-threaded comparison, TEDDY showed lower runtime and peak memory usage than ChimeraTE while retaining isoform-structure-level resolution. For larger mammalian or multi-sample datasets, TEDDY can additionally use thread-level parallelization during reconstruction and sample-level parallelization across independent inputs.

**Flow Cytometry Gating Strategy**

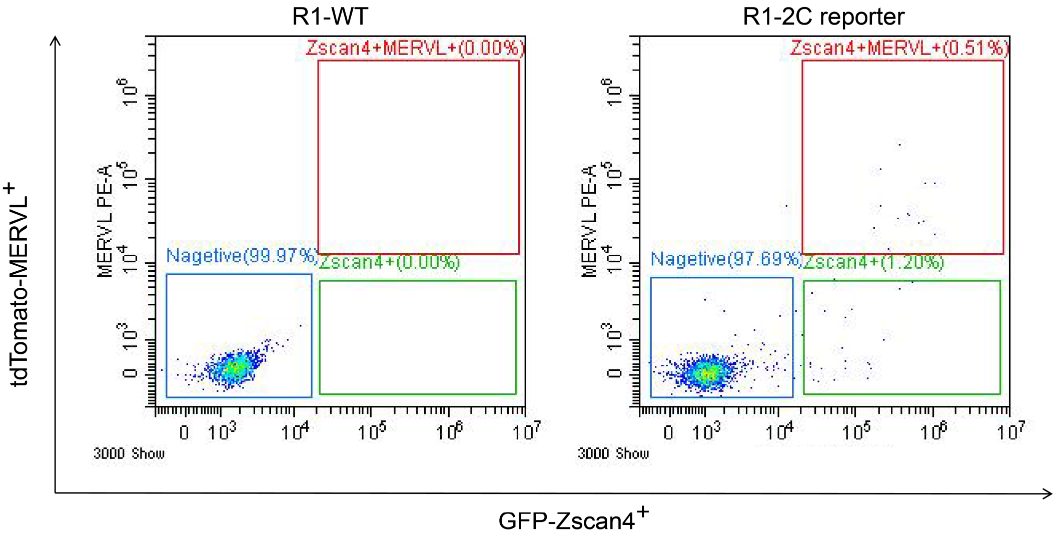


Flow cytometric gating strategy for the isolation of MERVL$^-$/*Zscan4*$^+$ and MERVL$^+$/*Zscan4*$^+$ cells. Cells were initially gated based on forward and side scatter to exclude debris, followed by doublet discrimination to retain singlets. The gates defining MERVL–tdTomato and *Zscan4*–EGFP positivity were established using the corresponding negative-control cells. MERVL$^-$/*Zscan4*$^+$ and MERVL$^+$/*Zscan4*$^+$ populations were subsequently sorted, whereas the unsorted parental ESC population was used as the control.
